# Type 1 LacNAc and LacdiNAc on *N*-glycans as molecular biomarkers of human melanoma cells

**DOI:** 10.1101/2025.10.27.684785

**Authors:** Karolina Grzesik, Paweł Link-Lenczowski, Andrea Carpentieri, Angela Amoresano, Manfred Wuhrer, Dorota Hoja-Łukowicz

## Abstract

Melanoma, the most dangerous form of skin cancer, is characterized by its high potential to spread or metastasize. Tumor progression is associated with changes in glycosylation that occur early in carcinogenesis and evolve as the cancer develops and spreads. To identify new melanoma transformation markers, we analyzed N-glycans from melanocytes and melanoma cell lines at different progression stages. We performed analyses using matrix-assisted laser desorption/ionization (MALDI)-mass spectrometry (MS) and hydrophilic interaction liquid chromatography (HILIC)-high performance liquid chromatography (HPLC) techniques on N-glycans without and after digestion with exoglycosidase arrays. Our results showed that, unlike melanocytes, melanoma cells express higher levels of type 1 LacNAc units and triantennary complex type glycans instead of high-mannose type glycans. Our results revealed that the N-glycomes of all analyzed cell lines possess Lewis X/A epitopes and, for the first time, demonstrate the presence of Galβ1-4Galβ1-4GlcNAc and Galβ1-3Galβ1-3GlcNAc units. The characteristic feature of melanoma cells is the presence of LacdiNAc structures. Our study provides the first comprehensive characterization of the N-glycome of melanocytes and suggests novel glyco-biomarkers of melanoma progression.

## Introduction

Melanoma develops from melanocytes, which are located primarily in the epidermis, the outermost layer of the skin. The American Joint Committee on Cancer (AJCC) melanoma staging system considers T/N/M category, tumor thickness, ulceration, and mitotic rate (Gershenwald et al., 2017). This system is widely used for melanoma classification, prognostic prediction, and individualized therapy planning. Lactate dehydrogenase (LDH) serum levels are the only molecular biomarker used to stratify patients into AJCC stages (Keung and Gershenwald, 2018). Detecting and surgically removing melanoma lesions at an early stage is associated with a 90% likelihood of cure. However, when melanoma has spread to other organs, the patient’s chances of survival decrease drastically, and treatment with chemotherapy, radiotherapy, immunotherapy, or targeted therapies becomes ineffective (Kenneth, 2018). The median survival after distant metastasis is only six to nine months, and the five-year survival rate is less than five percent (Matthews et al., 2010). Therefore, early disease detection and timely treatment initiation are crucial. Melanoma remains a cancer for which diagnostic and prognostic markers, as well as new therapeutic targets, are necessary.

Many studies have shown that alterations in protein glycosylation occur during oncogenic transformation. These changes impact cancer cell growth, survival, invasion, and metastasis (Stowell et al., 2015, Hoja-Łukowicz et al., 2017, Grzesik and Hoja-łukowicz, 2021). Tumor-associated carbohydrate antigens can serve as potent prognostic and diagnostic markers, as well as therapeutic targets (Grzesik et al., 2023). Although studies describing changes in the *N*-glycosylation profile of proteins during melanoma progression exist, this issue remains largely unexplored. Furthermore, most of these studies have examined individual glycoproteins (Ciolczyk-Wierzbicka et al., 2002, Pocheć et al., 2015, Hoja-Łukowicz et al., 2013, Abrahams, 2016, Agrawal et al., 2017). Virág et al. (2021) proposed that changes in the fucosylation and α2,6-sialylation of serum A1AG could serve as diagnostic markers for distinguishing high-risk melanoma patients from healthy individuals. In the present study, we determined and compared the total *N*-glycan profiles released from HEMa-LP (melanocytes), WM793 (primary skin melanoma; vertical growth phase), and WM266-4 (metastatic melanoma). Based on our results, we propose the LacNAc type 1 epitope as a new glycan marker of melanoma progression.

## Materials and methods

### Materials

RPMI 1640 with GlutaMax-I medium, Medium 254 and Human Melanocyte Growth Supplement (HMGS-2 PMA-free), Penicillin-Streptomycin solution, Gibco Fetal bovine serum and Formic acid solution 50%, Honeywell Fluka were purchased from Thermo Fisher Scientific (USA). Supelco Supelclean™ ENVI-Carb™ SPE Tube, 2-aminobenzamide (2-AB), sodium cyanoborohydride, α2-3-neuraminidase from *Streptococcus pneumonia*, neuraminidase from *Arthrobacter ureafaciens*, α-galactosidase from green coffee beans, *Streptococcus pneumoniae* hexosaminidase recombinant in *E. coli* and N-glycosidase F (PNGase F) from *Elizabethkingia meningoseptica* were purchased from Sigma-Aldrich (USA). β-galactosidase from *Streptococcus pneumoniae*, β-galactosidase from bovine testes, β-N-acetylhexosaminidase from jack bean, α-fucosidase from bovine kidney, α-mannosidase from jack bean and β-mannosidase from *Helix pomatia* were obtained from Prozyme (USA). β1-3-galactosidase from *Xanthomonas manihotis* was purchased from New England Biolabs, α1-3,4 Fucosidase Kit (fucosidase from *Ruminococcus gnavus*), α1-3,4 fucosidase from *Xanthomonas manihotis* and 2-AB Glucose Homopolymer Ladder were from Ludger (UK). J.T.Baker Acetonitrile, HPLC Far UV/Gradient grade was obtained from S.Witko (Poland). Ammonium acetate for HPLC was from POCH (Poland). MycoAlert^TM^ mycoplasma detection kit was from Lonza. All other chemicals were of the highest purity and were purchased from Sigma (USA).

### Cell lines

Normal human primary epidermal melanocytes HEMa-LP were obtained from Thermo Fisher Scientific (USA). Human cutaneous primary melanoma cell line WM793 from the vertical growth phase (VGP) and metastatic melanoma cell line WM266-4 (lymph-node metastasis) were obtained from the ESTDAB Melanoma Cell Bank (Tübingen).

### Cell culture conditions and cell extract preparation

HEMa-LP cells were maintained in 254 medium with HMGS-2 PMA-free. WM266-4 and WM793 cells were maintained in RPMI 1640 medium with GlutaMax-I supplemented with 10 % fetal bovine serum, 100 units/ml penicillin and 100 μg/ml streptomycin. Cells were grown in monolayers in a 5 % CO_2_ atmosphere at 37 °C in a humidified incubator. After reaching 80-90 % confluence, the cells were washed 3×4 ml DPBS and harvested. Cell cultures were conducted three times performing the entire experiment in triplicate. All cell cultures were free of mycoplasma infection as verified by MycoAlert^TM^ mycoplasma detection kit.

### Release and purification of oligosaccharides

Five miligrams of lyophilized cell pellets were solubilized and denatured in 300 µl 0.5% SDS and 2% β-mercaptoethanol at 100°C for 10 min. One hundred and fifty microliters of 150 mM NH_4_HCO_3_, pH 7.8 and 55 µl Nonidet P40 (1%) were added and oligosaccharides were cleaved from proteins by digestion with 2.5 unit of N-glycosidase F for 24 h at 37°C. Thereafter, 2 ml of cold ethanol was added, and the samples were centrifuged for 20 min at 2600x g at 4°C. The supernatants were then spun again for 30 min at 20 000x g at 4°C. Oligosaccharides were purified on SupelClean™ ENVI-Carb™ SPE Tube columns according to the manufacturer’s protocol and then dried in vacuo.

### Fluorescent labeling of the oligosaccharides

The oligosaccharides were labeled with 2-AB via reductive amination for three hours at 65 °C, according to the method described by Bigge et al. (1995). The labeled oligosaccharides were then purified using octanal, following the method described by Chu et al. (2018).

### HILIC HPLC (Protocol 1)

The 2-AB-labeled oligosaccharides were dissolved in 200 µL of Milli-Q water. Aliquots of 1 µL from each sample were treated with different mixtures of exoglycosidases in 20 mM sodium acetate buffer (pH 5.5) overnight at 37 °C. The enzymes used and their specificity are presented in Table 1. Each reaction mixture was cleared of enzymes using an Amicon Ultra centrifugal filter with a 10-kDa cutoff membrane according to the manufacturer’s instructions. Then, the mixture was dried and frozen at −30 °C.

**Table 1.**
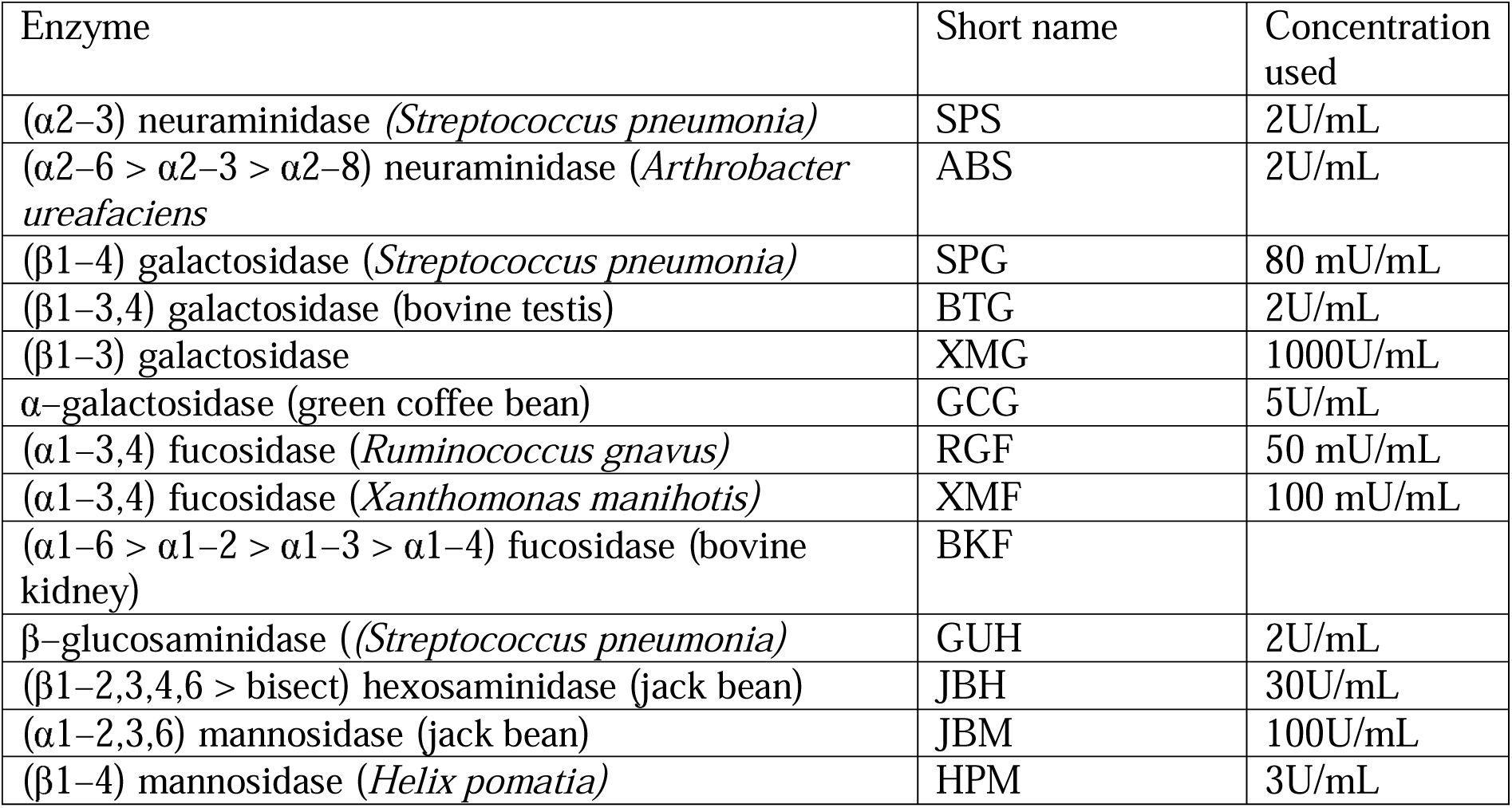
List of enzymes used.

| Enzyme | Short name | Concentration used |
| --- | --- | --- |
| (α2–3) neuraminidase (*Streptococcus pneumonia*) | SPS | 2U/mL |
| (α2–6 > α2–3 > α2–8) neuraminidase (*Arthrobacter ureafaciens*) | ABS | 2U/mL |
| (β1–4) galactosidase (*Streptococcus pneumonia*) | SPG | 80 mU/mL |
| (β1–3,4) galactosidase (bovine testis) | BTG | 2U/mL |
| (β1–3) galactosidase | XMG | 1000U/mL |
| α–galactosidase (green coffee bean) | GCG | 5U/mL |
| (α1–3,4) fucosidase (*Ruminococcus gnavus*) | RGF | 50 mU/mL |
| (α1–3,4) fucosidase (*Xanthomonas manihotis*) | XMF | 100 mU/mL |
| (α1–6 > α1–2 > α1–3 > α1–4) fucosidase (bovine kidney) | BKF | |
| β–glucosaminidase ((*Streptococcus pneumonia*) | GUH | 2U/mL |
| (β1–2,3,4,6 > bisect) hexosaminidase (jack bean) | JBH | 30U/mL |
| (α1–2,3,6) mannosidase (jack bean) | JBM | 100U/mL |
| (β1–4) mannosidase (*Helix pomatia*) | HPM | 3U/mL |

The undigested and exoglycosidase-digested glycan samples were dissolved in 50 µL of 80% acetonitrile. They were then separated using a Shimadzu Prominence HPLC system with two Shimadzu LC-20AD pumps and a Shimadzu RF-20Axs fluorescence detector (Japan; λ_(ex)=330 nm, λ_(em)=420 nm). The samples were separated on a 4.6 × 250 mm TSKgel Amide-80 column maintained at 30 °C. The gradient used is as follows: Solvent A was acetonitrile and Solvent B was 50 mM ammonium formate at pH 4.4. Initially, the conditions were 20% B at a flow rate of 0.4 mL/min. Then, there was a linear gradient of 20–53% B over 132 min, followed by 53–100% B over the next 3 min. The column was then washed with 100% B at a flow rate of 1 mL/min for five minutes before reequilibrating in the initial solvent system. The 2-AB glucose homopolymer ladder was used as an external standard. A well-established fifth-order polynomial relationship was used to predict the glucose unit (GU) value based on retention time.

### LC-MS (Protocol 2)

The 2-AB-labeled *N*-glycans were analyzed using ultra-performance liquid chromatography (UPLC) with fluorescence detection coupled to ion mobility spectrometry–quadrupole time-of-flight mass spectrometry (IMS–qToF MS) via an electrospray ionization (ESI) source (ACQUITY I-Class Plus UPLC and Vion IMS–qToF, Waters). Chromatographic separation was performed on a hydrophilic interaction liquid chromatography (HILIC) column (ACQUITY UPLC Glycan BEH Amide, 130 Å, 1.7 µm, 2.1 × 150 mm; Waters) maintained at 65 °C. The mobile phases consisted of 50 mM ammonium formate, pH 4.4 (solvent A), and acetonitrile (solvent B). The gradient program was as follows: t = 0.0 min, 25% A at 0.4 mL/min; t = 35.0 min, 46% A at 0.4 mL/min; t = 36.0 min, 100% A at 0.2 mL/min; t = 39.5 min, 100% A at 0.2 mL/min; t = 43.5 min, 25% A at 0.2 mL/min; t = 47.6 min, 25% A at 0.4 mL/min; t = 55.0 min, 25% A at 0.4 mL/min. Fluorescence detection was carried out with excitation and emission wavelengths set at 330 nm and 420 nm, respectively. Mass spectrometric analysis was performed in positive ion time-of-flight (ToF) MS mode, acquiring ions in the m/z range 600–2000. Mass accuracy was maintained by sampling a reference mass standard every 60 s. Instrument parameters were set as follows: capillary voltage, 3.0 kV; source temperature, 120 °C; desolvation temperature, 350 °C; desolvation gas flow, 800 L/h.

### MALDI-TOF MS (Protocol 3)

Mass spectrometry analysis was performed using an AB SCIEX 5800 MALDI-TOF mass spectrometer with Voyager software.

The 2-AB-labeled *N*-linked glycans were resuspended in a 1:1 (v/v) solution of water and methanol. Then, 0.5 µL volumes of the samples were mixed with an equal volume of matrix solution consisting of 10 mg/mL 2,5-dihydroxybenzoic acid (DHB) in a 1:1 (v/v) solution of water and methanol with 0.2% HCOOH on a MALDI metal plate. MS spectra were acquired in positive reflection ion mode. Mass calibration was performed using external peptide standards purchased from Applied Biosystems. Spectra were acquired using a mass-to-charge (m/z) range of 800–4,000. Analysis of each sample involved the use of the same laser intensity and number of shots across the entire sample plate spot to ensure homogeneous analysis of the analyte of interest. Raw data were analyzed using the mMass-Open Source Mass Spectrometry Tool program. Glycans were analyzed by manually inspecting queries in the GlycoWorkbench program.

### MALDI-TOF MS (Protocol 4)

2-AB-labeled *N*-glycan samples were subjected to sialic acid linkage-specific derivatization, resulting in the ethyl esterification of α2,6NeuAc (derivatized mass: 319.127 Da) and the amidation of α2,3NeuAc (derivatized mass: 290.111 Da). For this, 2-AB-labeled *N*-glycans were incubated with 20 µL ethyl esterification reagent (250 mM EDC and 250 mM HOBt in ethanol) for 30 min at 37°C. The reaction mixture was further incubated with 4 µL 28% NH_4_OH for 30 min at 37°C for the conversion of α2,3-linked sialic acids into amide derivatives (Lageveen-Kammeijer 2019). Subsequently, *N*-glycans were purified by cotton hydrophilic interaction liquid chromatography solid-phase extraction (HILIC SPE)(Vreeker 2018). Samples were eluted in 10 µL of deionized water, and 2 µL of sample were spotted together with 1 µL sDHB matrix (5 mg/mL in 99% ACN with 1 mmol/L NaOH) onto a polished-steel MALDI target plate. Samples were measured in reflectron-positive mode on a rapifleX MALDI-TOF MS (Bruker Daltonics, Bremen, Germany) and data were processed as described for Protocol 3.

## Results

The aim of the study was to identify novel glycomarkers of human cutaneous melanoma by determining and comparing the *N*-glycome of melanocytes (HEMa-LP cell lines) with the *N*-glycome of cutaneous melanoma cells at different stages of cancer development, such as the vertical growth phase (WM793 cell line) and lymph node metastasis (WM266-4 cell line). N-glycans were released from glycoproteins using PNGase F, labeled with 2-AB, and analyzed in four laboratories using different protocols. In protocol 1, the HILIC HPLC method was used to separate the *N*-glycans before and after digesting them with different exoglycosidase arrays. Each post-reaction mixture was fractionated separately, and the glycan structures were deduced using the GlycoStore database, taking into account the changes in peak positions described by GU values, as well as the changes in relative peak areas after treatment with the given exoglycosidase mixture. The *N*-glycome of all cell lines was analyzed in triplicate. The results of this experiment can be found in Supplementary Tables S1–S3. Supplementary Figure S1 shows selected chromatograms for a set of digestion arrays using *Streptococcus pneumoniae* β1-4 galactosidase (SPG). In protocol 2, the UPLC-MS method was used to analyze undigested *N*-glycan pools. The *N*-glycan structures were identified for each chromatographic peak based on GU values and confirmed by accurate mass measurements. GU values and mass data were then compared to reference databases using the Waters UNIFI Scientific Information System to assign glycan structures. For the quantitative analysis, the fluorescence peak areas were integrated and the relative abundance of each glycan was expressed as a percentage of the total glycan pool. This analysis was carried out in triplicate. The results of this experiment are presented in Supplementary Table S4. The resulting chromatograms are presented in Supplementary Figures S2–S4. Protocol 3 is based on MALDI-TOF-MS analysis of desialylated *N*-glycans. Using the GlycoWorkbench program (https://glycoworkbench.software.informer.com/2.1/) and experimentally determined masses, all possible glycan structure compositions were proposed. This analysis was performed once, and the results are presented in Supplementary Table S5. In protocol 4, *N*-glycans underwent ethyl esterification and amidation (EEA) of sialic acid residues or digestion with a mixture of *Arthrobacter ureafaciens* sialidase (ABS) and SPG, followed by MALDI-TOF-MS analysis. The obtained mass spectra are shown in Supplementary Data 1. They can be viewed using the mMass program. Peak lists for spectra with m/z, intensity, and signal-to-noise (s/n) ratio are presented in Supplementary Data 2. The GlycoWorkbench program was used to propose all possible compositions of glycan structures with linkage-specific differentiation of sialic acids or the presence of β1-3-linked galactose residues on the basis of experimentally determined masses. These analyses were carried out in triplicate, and the results are presented in Supplementary Table S6. Table 2 lists all the *N*-glycans identified in this study for each cell line. This table is a compilation of structures obtained from analyses using HILIC HPLC (protocol 1), UPLC-MS (protocol 2), and MALDI-TOF-MS after EEA derivatization of sialic acids (protocol 4). ABS-treated and HILIC HPLC-profiled oligosaccharides were confirmed by MALDI-TOF-MS analysis of desialylated *N*-glycans (protocol 3). Conversely, the products of ABS+SPG digestion detected by HILIC HPLC were confirmed by MALDI-TOF-MS analysis (protocol 4). These confirmations are included in Supplementary Tables 1–3 (columns H–J, V–W, and BM–BN).

**Table 2.**
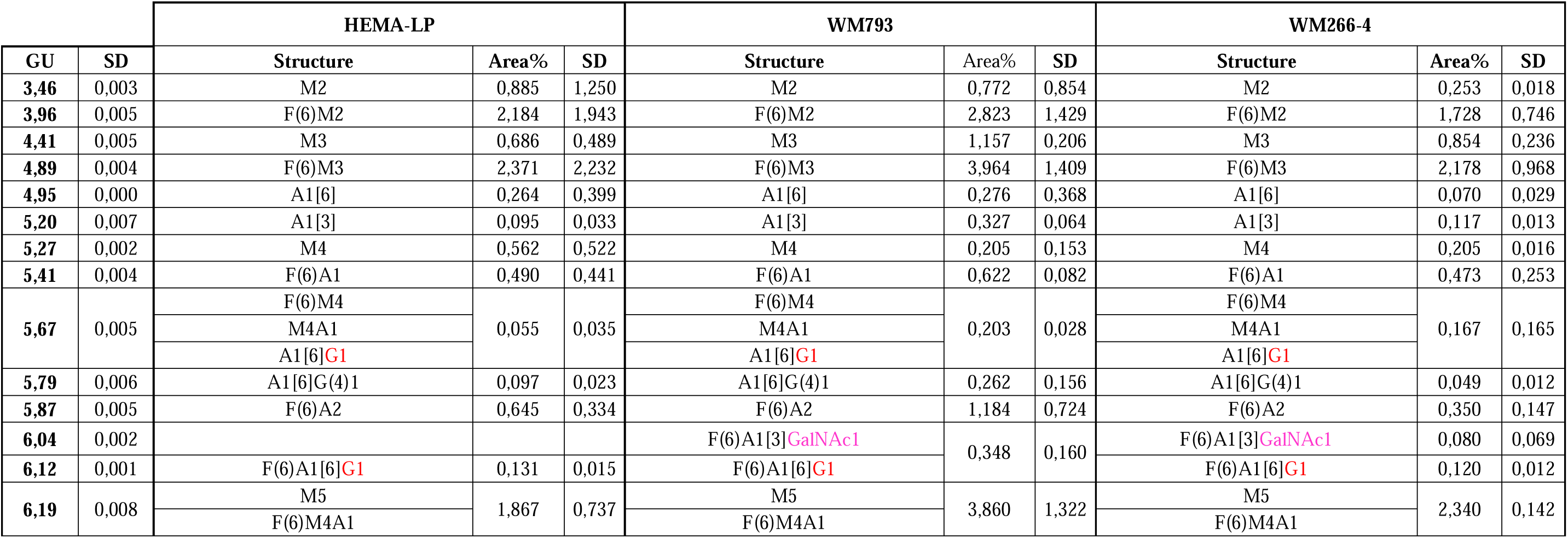
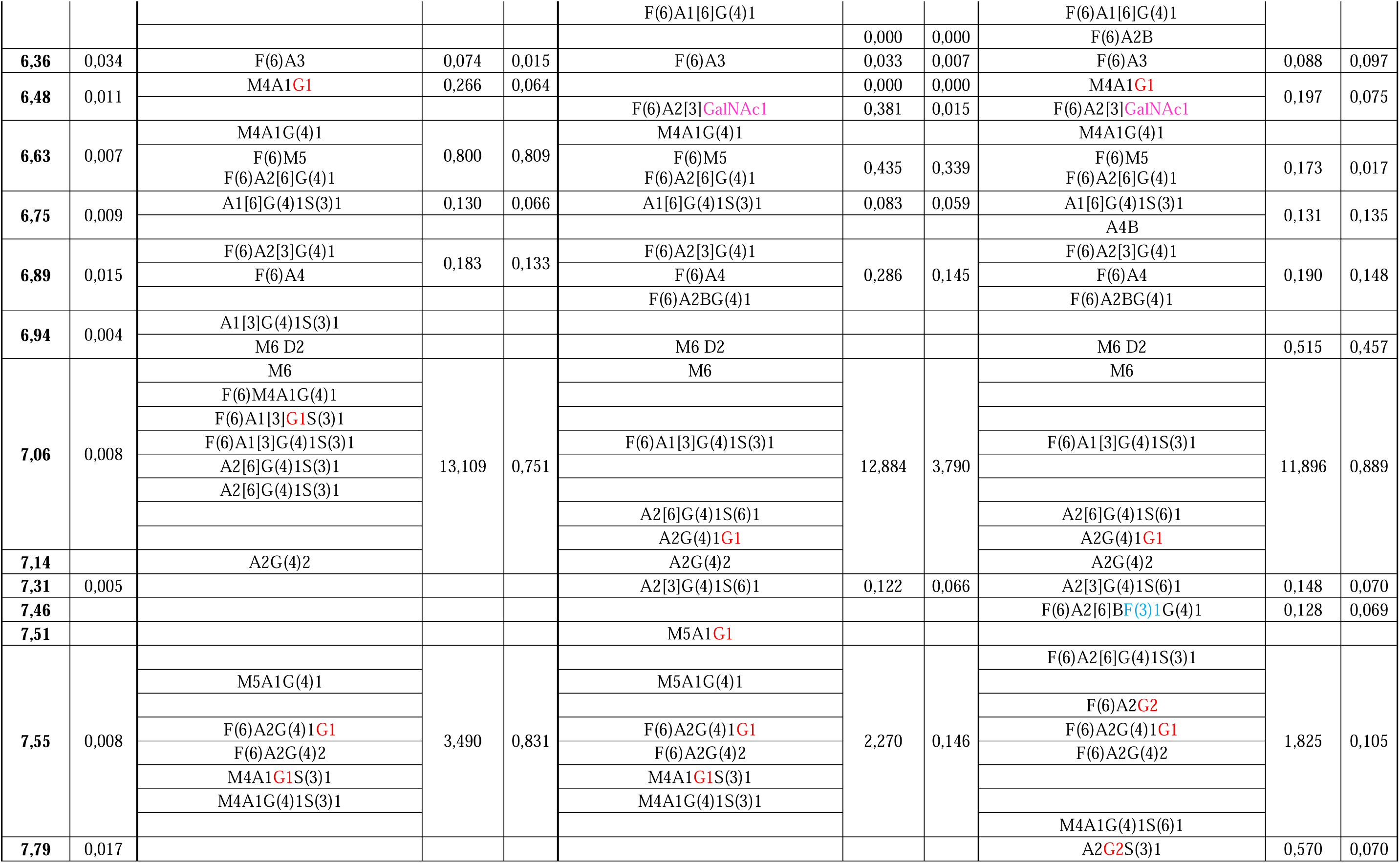
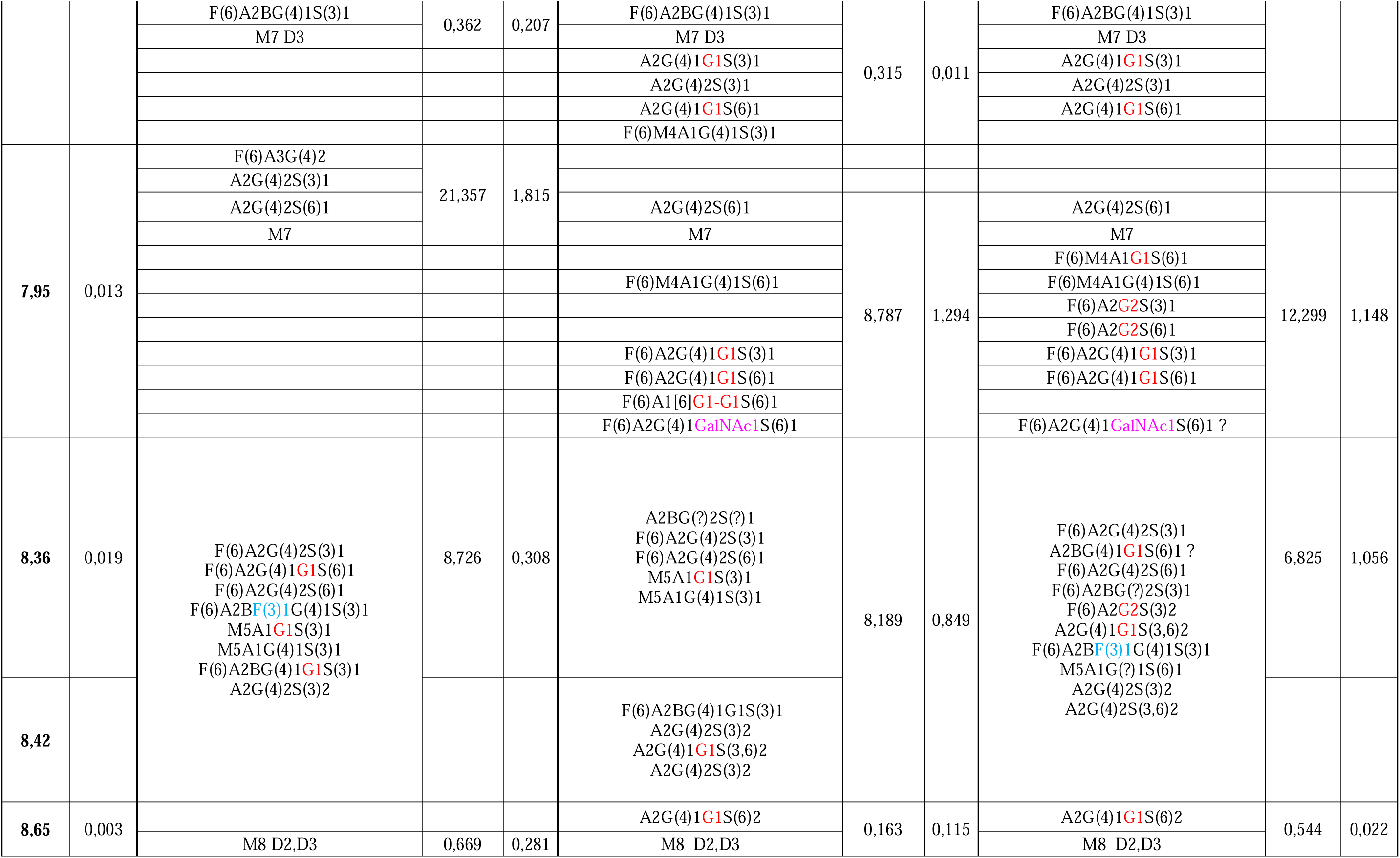
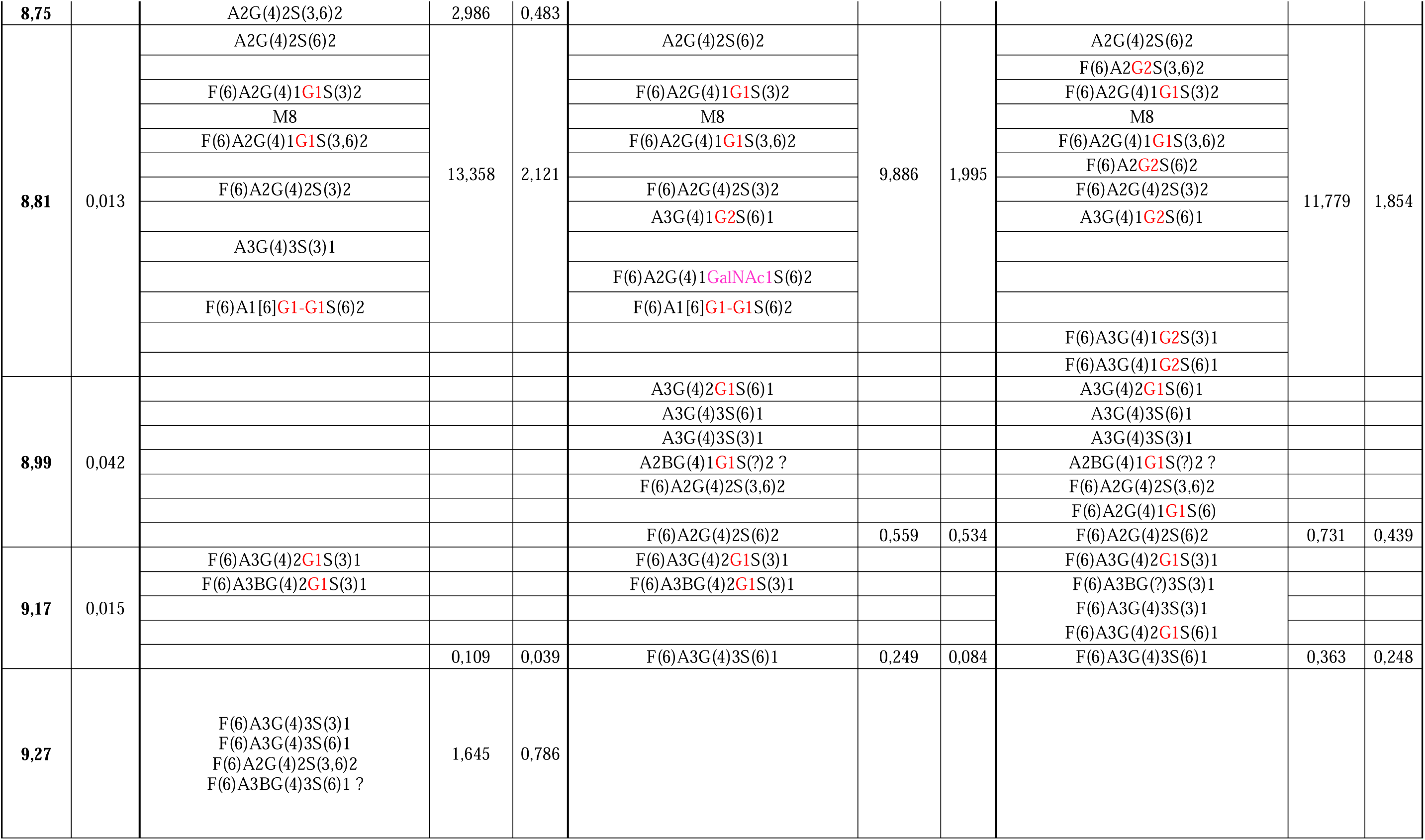
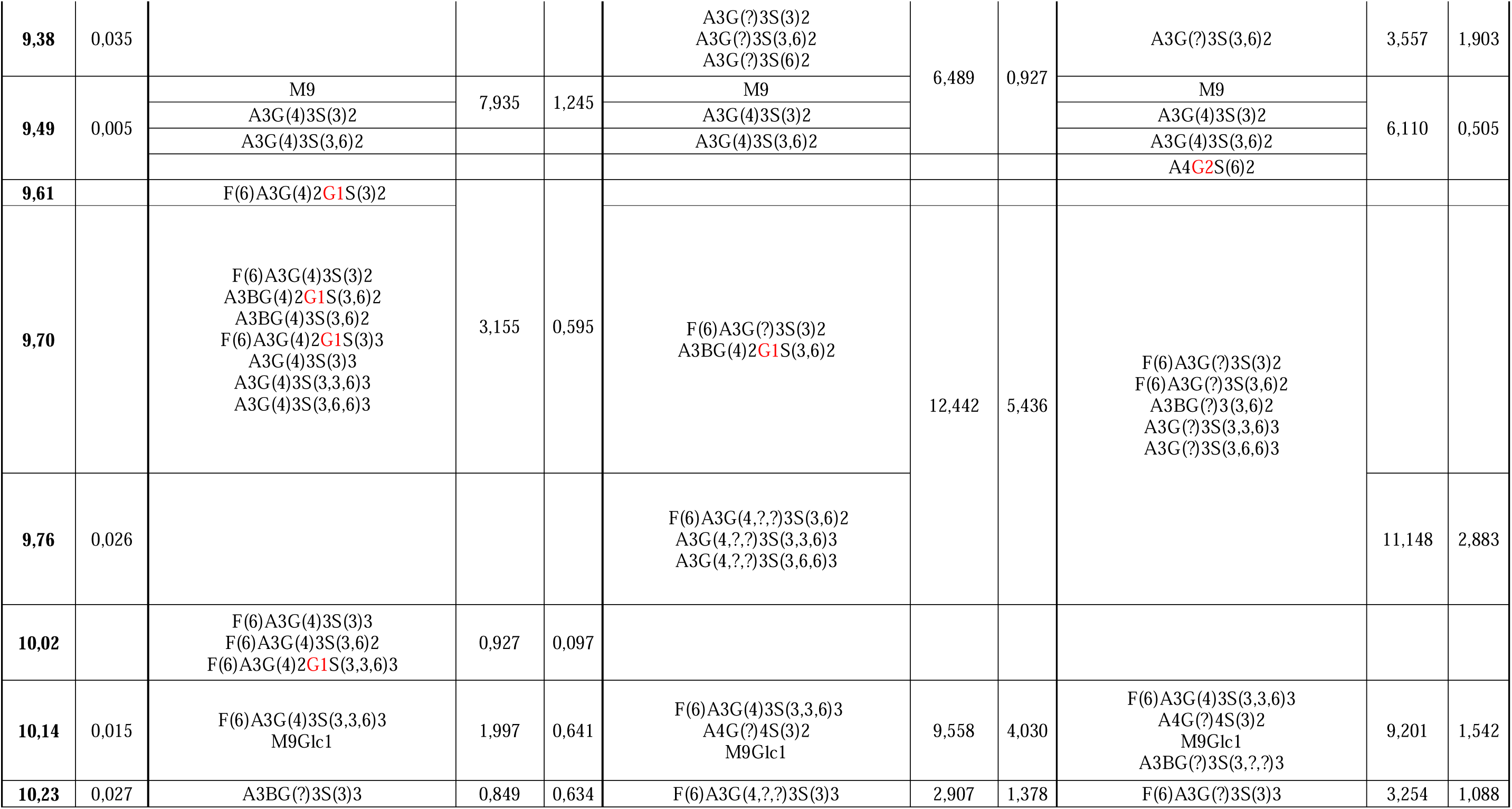
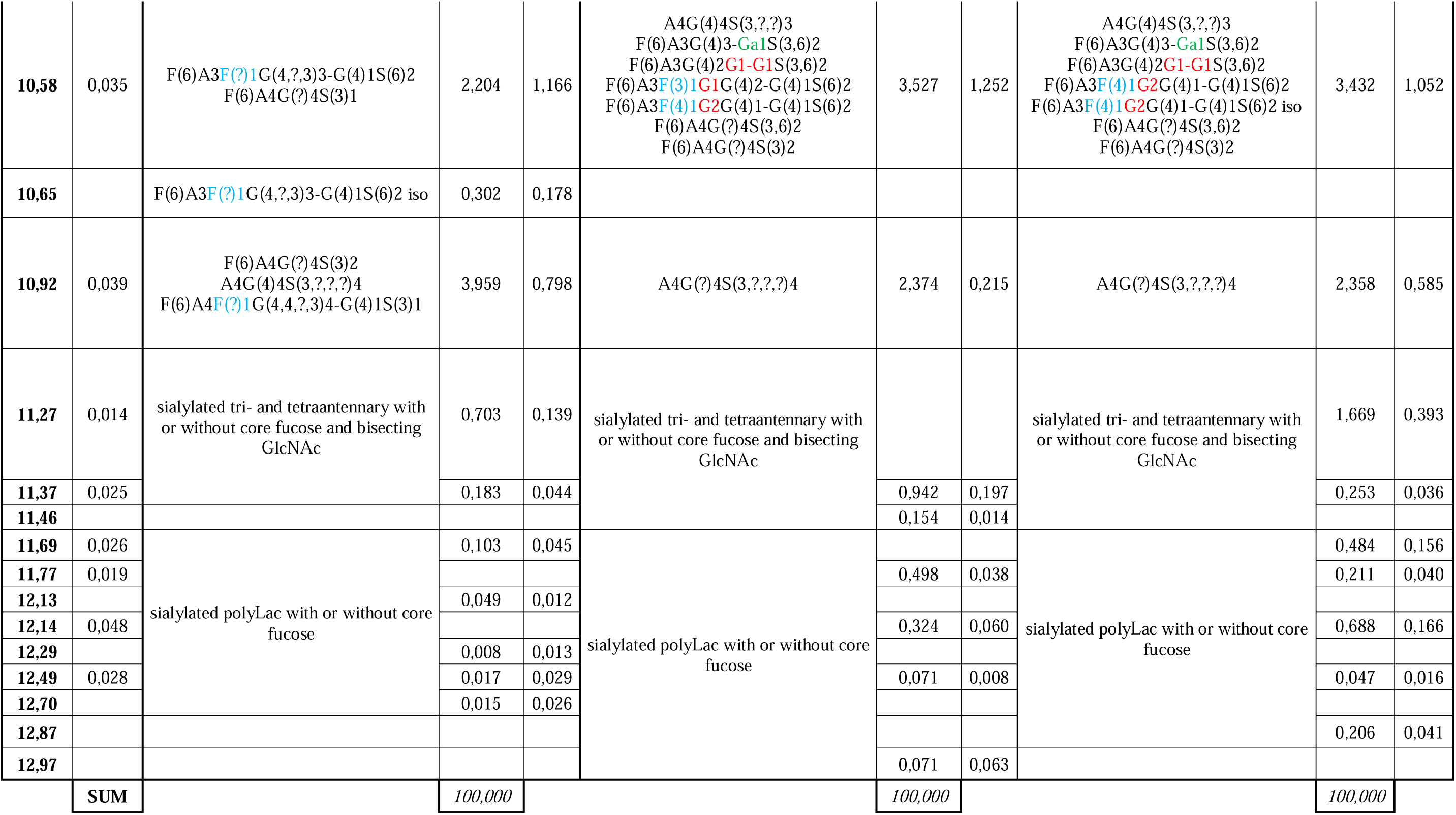
*N*-Glycans of melanocytes (HEMA-LP), primary melanoma (WM793), and metastatic melanoma (WM266-4) cell lines identified by HILIC HPLC. Relative percent abundance was calculated based on the area of the chromatographic peaks, expressed as a percentage of the total glycan abundance. The mean relative abundance (Area%) and standard deviation (SD) are shown for n = 3. The glucose unit (GU) value of each peak was calculated by comparing it to the dextran hydrolysate ladder separation. The structures were assigned based on the glucose unit values, the known incremental values for monosaccharide residues, and the known specificity of the exoglycosidase enzymes. The structure abbreviations used are as follows: Aa[3/6] indicates the number "a" of antennae on the trimannosyl core linked to the 3/6-mannose arm. G(4)b and G(c) indicate the number "b" and "c" of terminal galactose residues, respectively, linked β1-4 and β1-3 to antenna GlcNAc. F(6) and F(3/4)d indicate a core fucose, respectively, linked α1-6 to the core GlcNAc and "d" fucose residues, respectively, linked α1-3/4 to antenna GlcNAc. B represents a bisecting GlcNAc, respectively, linked β1-4 to core mannose. Me represents the number "e" of mannose residues. S(3/6)f represents the number "f" of sialic acids α2-3/6-linked to galactose or *N*-acetylhexosamine in antennae.

### Sialic acid linkage-isomers

After treatment with *A. ureafaciens* sialidase (ABS), which has specificity for α2-3,6,8-linked terminal NeuAc and NeuGc, *N*-oligosaccharides from all analyzed cell lines were separated into five dominant peaks in HILIC HPLC analysis. The GU values of these peaks were in the range of 7.04-7.07, 7.48-7.51, 7.92-7.95, 8.78-8.82, and 9.45-9.55. For melanoma cells, an additional dominant peak with a GU value ranging from 8.25 to 8.28 was observed. This peak occupied approximately 17% and 18% of the WM793 and WM266-4 *N*-glycomes, respectively; however, it did not exceed 2% in melanocytes. The GU value of this peak corresponds to the tri-galactosylated tri-antennary sugar structure. For HEMA-LP cell lines, S. pneumoniae sialidase digestion (SPS, which removes α2-3-linked sialic acids) produced distributions and surface areas of peaks that were very similar to those obtained after ABS digestion. This suggests that melanocyte-derived *N*-glycans mainly contain α2-3-linked sialic acids. Generally, no more than one α2-6-linked sialic acid residue per structure was observed for mixed types of sialic acid bonding in melanocytes. Exceptions were isomeric structures with a molecular mass of 3162 Da and detected in peaks of 10.51 and 10.63 GU by UPLC-MS (Supplementary Table S4). The EEA derivatization method confirmed these structures as containing two α2-6-linked sialic acid residues (*m/z* 3241; Supplementary Table S6).

In contrast, digesting melanoma-derived *N*-oligosaccharides with SPS produced new peaks not present in chromatograms after ABS digestion. For instance, new peaks with GU values of 8.95–9.01 and 9.23–9.27 occupied 6.6% and 1%, respectively, of the WM793 cell *N*-glycome, and 7.8% and 1.7%, respectively, of the WM266-4 cell *N*-glycome. These peaks correspond to *N*-glycans containing at least one α2-6-linked sialic acid residue. Another peak, specific to melanoma samples and resulting from SPS digestion, contained tri-galactosylated tri-antennary oligosaccharides with two α2,6-linked sialic acid residues and had a GU value in the range of 9.74-9.78. This peak accounted for over 2.2% and 2.9% of the *N*-glycans in WM793 and WM266-4 cells, respectively. The conclusions drawn from the HILIC HPLC method, were confirmed, by MALDI-MS analysis of the EEA derivatization products. Based on Protocol 4 results, melanoma cells appear to be more sialylated than melanocytes. Melanocytes are generally characterized by a higher percentage of mono α2,3-sialylated structures compared to melanoma cells. Conversely, melanoma cells have more glycans with α2,6-linked sialic acid in mono-, di-, or trisialylation forms per structure. Figure 1 shows the quantitative analysis that supports these conclusions, based on the data in Supplementary Tables S6-S7, and Supplementary Data 2.

**Figure 1.**
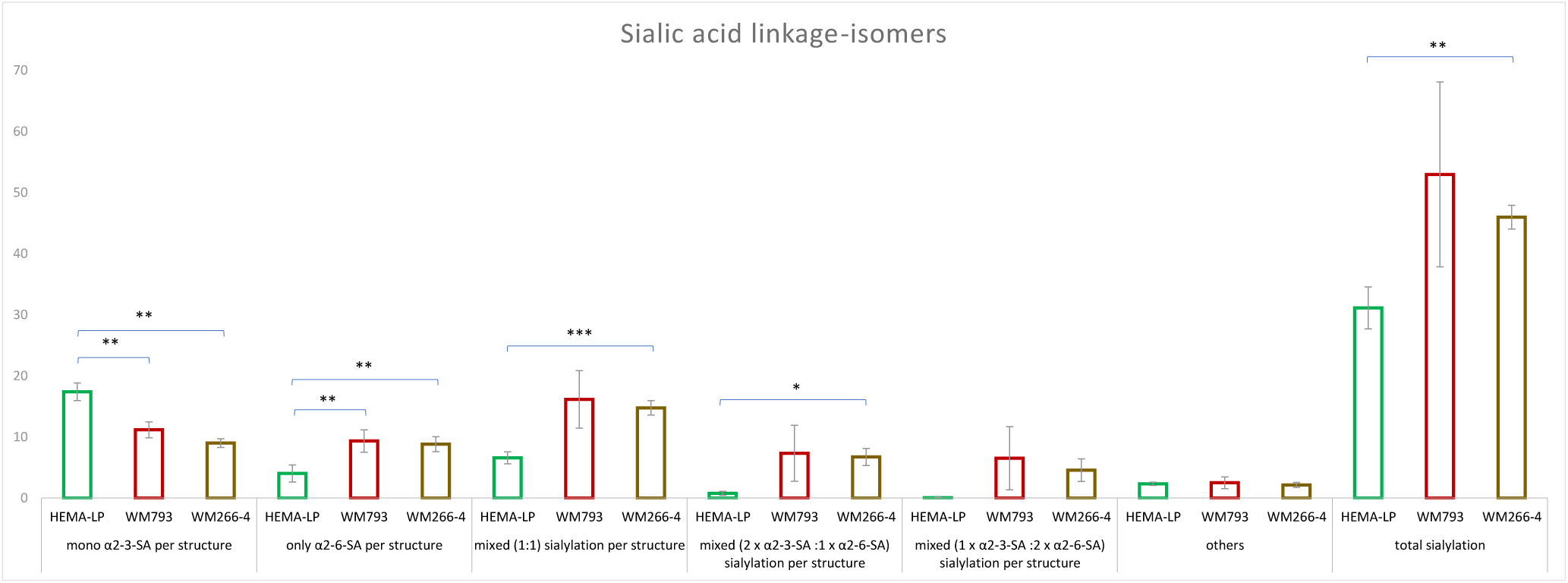
Relative percentage abundance of sialic linkage-isomers per structure detected by MALDI-TOF MS after EEA derivatization of sialic acids. S(3) = α2-3 linked sialic acid, S(6) = α2-6 linked sialic acid. The corresponding data are presented in Supplementary Table S7. Error bars represent standard deviation (n = 3). * p < 0.05; ** p < 0.01; *** p < 0.001 (t-test in Excel).

### LacNAc type 1 unit

To detect galactose linkage isomers, we performed sequential digestions with exoglycosidases, followed by HILIC HPLC separations. One set of exoglycosidase arrays contained S. pneumoniae β-galactosidase (SPG), which removes β1-4-bound galactose, and the other contained bovine testicular β-galactosidase (BTG), which removes β1-3/4-bound galactose. Digesting melanocyte-derived oligosaccharides with mixtures of ABS + SPG or ABS + BTG resulted in six main peaks with GU ranges of 5.43–5.44, 5.84–5.86, 7.03–7.05, 7.92–7.95, 8.78–8.81, and 9.45–9.48. The percentages of the main peaks were similar in both post-reaction mixtures, ranging from 8% to just over 16%. These peaks mainly contained A2, F(6)A2, and high-mannose M6-M9 oligosaccharides. Structures containing a single β1-3-linked galactose residue were assigned to corresponding peaks after ABS + SPG digestion and HILIC HPLC separation. These structures were of the hybrid type and of the complex type, with or without core fucose, and were located in peaks at GU values between 5.86 and 7.95 (see Supplementary Table S1). Ten out of sixteen structures were confirmed by MALDI-MS after ABS + SPG digestion and corresponded to the following m/z values: 1742, 1767, 1970, 2027, 2173, and 2376 (Supplementary Table S6). Digesting with ABS + SPG and jack bean β-N-acetylhexosaminidase (JBH), which has specificity for β1-2,3,4,6-linked GlcNAc and GalNAc, resulted in removing unsubstituted HexNAc residues from complex-type *N*-glycans. This process formed structures with single β1-3-linked galactose residues, which are mainly found on the α6 arm of *N*-glycans. Subsequent digestion with bovine kidney fucosidase (BKF), which cleaves core fucose more efficiently than other α-fucose linkages, moved these structures to new peaks with GU values between 5.64 and 5.87 (A1[6/3]G1). Further digestion with jack bean mannosidase (JBM; specificity for α1-2,3,6-linked mannose) resulted in the formation of some peaks corresponding to the M1 structure (2.63 GU) derived from pauci and high mannose type glycans, M4Glc1 structure (5.97 GU; and the incompletely digested M5Glc1 structure at 6.86 GU) derived from M9Glc1, and the M2A1[3/6]G1 structures (4.86 GU) derived from A1[3/6]G1 structures as well as from the hybrid type glycans M4A1G1 and M5A1G1. Trace amounts of two β1-3-linked galactose residues were found on structure with -GlcNAcβ1-3Galβ1-3Gal unit (assigned G1-G1), i.e. M2A1G1-G1 (5.65 GU), and on the structure carrying the Lewis A (LeA) epitope, i.e., M2A2F(4)1G2 (6.92 GU). These findings are discussed in the "Other structures" section.

In the case of *N*-oligosaccharides derived from melanoma WM793 and WM266-4 cells, digestion with a mixture of ABS + SPG and ABS + BTG enzymes gave a one dominant peak in the range of 5.84-5.86 GU after HILIC HPLC profiling (see Supplementary Tables S2-S3). This peak corresponded to A3 and F(6)A2 structures and occupied more than 30% and 39% and more than 25% and 30% of the total areas on chromatograms for the respective digestion products and cell lines. Furthermore, for both melanoma cell lines, HILIC HPLC profiles after ABS+SPG digestion showed a significantly higher peak at GU value of 6.60 compared to ABS+BTG digestion. This is due to the presence of a structure with a single β1-3-linked galactose residue (i.e. F(6)A2[6]G1), which corresponds to m/z 1767 in the **Supplementary Table S6**. This structure, as well as others containing a single β1-3-linked galactose residue, were identified from the m/z values obtained by MALDI-MS after ABS+SPG digestion (Supplementary Table S6) and assigned to the peaks in chromatograms (Supplementary Figure S1).

Unlike melanocytes, several *N*-glycans from melanoma cells were substituted with two β1-3-linked galactose residues. These residues were assigned to peaks with GU values between 7.41 and 8.81. The specific glycans were A3G2 (*m/z* 1986), A4G2 (*m/z* 2189), F(6)A4G2 (*m/z* 2335), and F(6)A4BG2 (*m/z* 2538). Additional structures specific to the WM266-4 metastatic melanoma cell line were also identified. These structures are A2G2 at 7.04 GU (*m/z* 1783), F(6)A2G2 at 7.56 GU (*m/z* 1929), F(6)A2BG2 at 7.62 GU (*m/z* 2132), A3BG2 and F(6)A3G2 at 7.94 GU (*m/z* 2189 and 2132 respectively) and F(6)A3BG2 at 8,27 GU (*m/z* 2335). Digestion with ABS + SPG + JBH removed the unsubstituted HexNAc residues and gave a dominant peak at 5.71-5.73 GU. This shows that the single β1-3-linked galactose is mainly located on the α6 arms of the *N*-glycans in both melanoma cell lines. This peak was enlaged as a result of digestion by the ABS + SPG + JBH + BKF mixture. Further digestion with JBM simplified all mannose-type glycans to the M1 and M4Glc1 structures (peaks with 2.64 GU and 5.95-5.97 GU; 6.83-6.88 GU for incomplete digested M5Glc1 structure), and shifted A1[6/3]G1, M4A1G1 and M5A1G1 structures to the peak with 4.85-4.86 GU. On the other hand, the A2G2 structure derived from tri- and tetra-antennary complex oligosaccharides lost a mannose residue and was shifted to the 6.08-6.10 GU peak (M2A2G2 structure). Part of the A2G2 structure that remained in the GU peaks between 6.90 and 7.01 originated from di-antennary structures (in the case of the WM266-4 cell line only) or from tri- or tetra-antennary structures in which one β1,3-linked galactose was present on the α3 arm and another on the α6 arm. Notably, the presence of M2A1[3/6]G1 structures at a peak of 4.86 GU was confirmed by their shift to a peak of approximately 4.06 GU after digestion with β1-3-galactosidase from *Xanthomonas manihotis* (XMG), as illustrated in Supplementary Figures S5–S7 and Supplementary Table S8-S10 (columns F–G). In all cell lines analyzed, enrichment of the exoglycosidase mixtures ABS + SPG + JBH + BKF + JBM or ABS + BTG + JBH + BKF + JBM with a β-mannosidase from *Helix pomatia* (HPM; specificity for β1-4-linked mannose) shifted the M1 structure to the peak corresponding to the GlcNAc2 structure (1.70 GU)(Supplementary Tables S1-3, Supplementary Figure S1). Finally, the percentage of oligosaccharides with a single β1-3-linked Gal residue was statistically significantly higher in melanoma WM793 and WM266-4 cells (17.01% and 18.88%, respectively) than in melanocytes (5.49%, p<0.001). Notably, the aforementioned content of a single β1-3-linked Gal residue in metastatic melanoma cells was also statistically significant compared to primary melanoma cells (p<0.05) (Figure 2, Supplementary Data 3).

**Figure 2.**
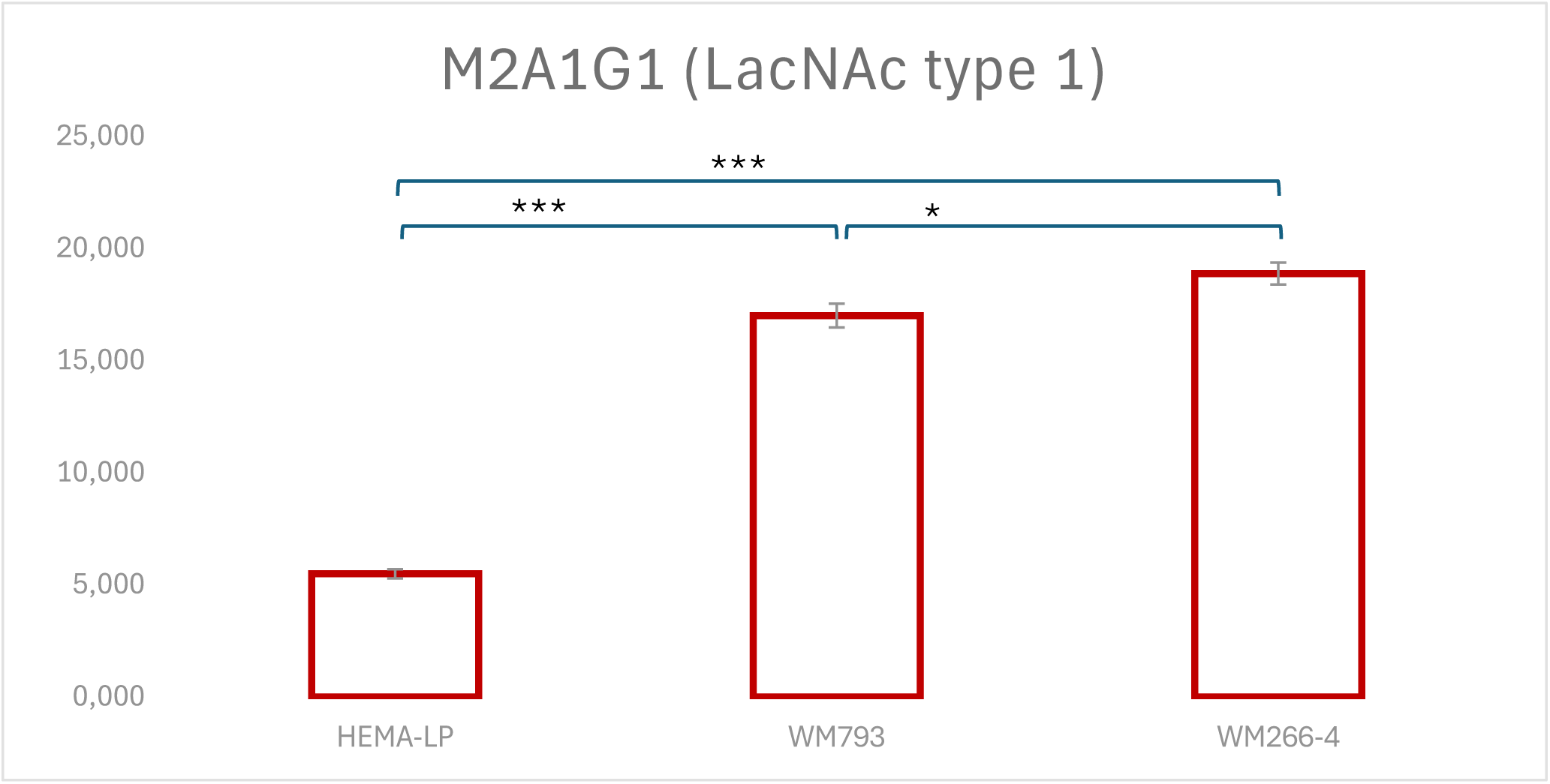
Relative percentage abundance of type 1 LacNAc identified by HILIC HPLC after digestion using the ABS+SPG+BKF+JBH+JBM assay. The enzymes used are ABS (α2-3,6,8 sialidase), BTG (β1-3,4 galactosidase), BKF (α1-6 fucosidase), JBH (β1–2,3,4,6 hexosaminidase), JBM (α1–2,3,6 mannosidase. The corresponding data are presented in Supplementary Data 3. Error bars represent standard deviation (n = 3). * p < 0.05; *** p < 0.001 (t-test in Excel).

#### LacdiNAc motif

To test for the presence of LacdiNAc structures, we performed a digestion of twice the amount of each sample with ABS, BTG, and *Streptococcus pneumoniae* hexosaminidase (GUH). GUH is specific to β-GlcNAc, but not to bisecting GlcNAc β(1-4)-linked to mannose. Then, we divided each sample into two equal parts and digested one part with JBH. For *N*-glycans from WM793 cells, we performed an additional test to detect LacdiNAc units. First, we digested twice the amount of the sample with a mixture of ABS, BTG, BKF, and GUH. After dividing the sample into two equal parts, we subjected one part to additional digestion with JBH. These digestions were performed once. We only identified *N*-glycan structures containing LacdiNAc in melanoma samples (see Figure 3 and Supplementary Tables S1 and S3 [columns BB-BG] and S2 [columns BB-BM]).

**Figure 3.**
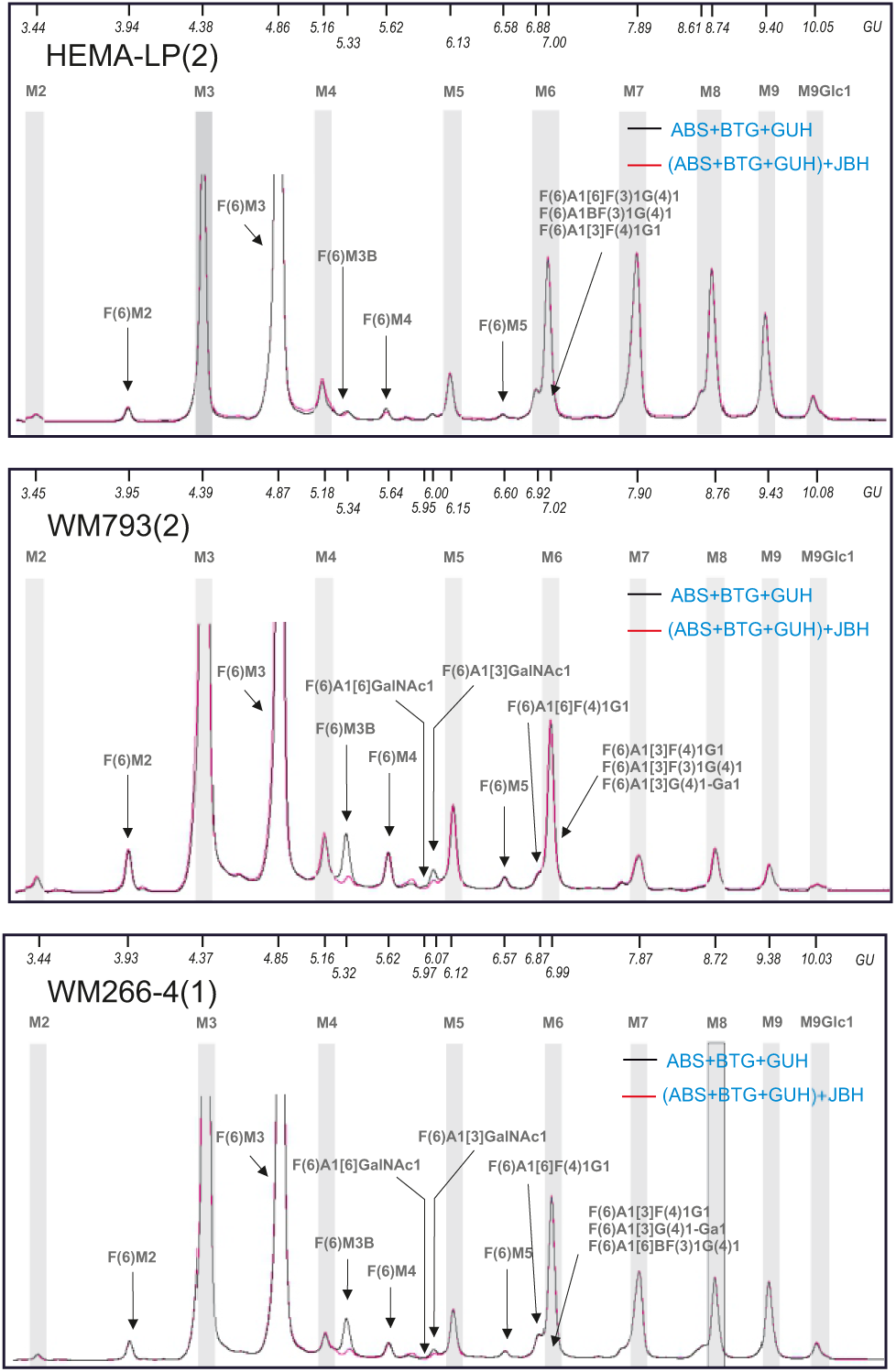
Identification of the LacdiNAc motif by HILIC-HPLC after exoglycosidase digestion. Twice the sample amount was used for digestion with ABS+BTG+GUH (α2-3,6,8 sialidase, β1-3,4 galactosidase, and β-N-acetylhexosaminidase). Each sample was then divided into two equal parts, with one part undergoing further digestion with β1–2,3,4,6 hexosaminidase (JBH). The corresponding data are presented in Supplementary Tables S1-S3 (columns BB-BG).

### *N*-glycan classes

Matrix-assisted laser desorption ionization mass spectrometry after EEA derivatization enabled us to determine the content of paucimannose and high-mannose-type *N*-glycans, as well as bi-, tri-, and tetra-antennary complex-type *N*-glycans. Figure 4 shows the percentage of each *N*-glycan class, calculated as the mean of three repetitions from the HEMa-LP, WM793, and WM266-4 cell lines. Over 65% of the oligosaccharides isolated from melanocytes were high-mannose *N*-glycans (M5 to M9). This result was statistically significant compared to the total amount of high-mannose-type glycans in metastatic melanoma cell lines (p < 0.05)(see Supplementary Data 4). An analysis of the percentage of individual high-mannose structures revealed that melanocytes have significantly higher amounts of M7 and M8 structures than primary melanoma cells (p < 0.005 and p < 0.02, respectively) and metastatic cells (p < 0.02 and p < 0.0002, respectively). Conversely, the M9GlcNAc1 structure was statistically significantly less prevalent in primary melanoma cells than in melanocytes and metastatic cells. The latter cell lines had a similar level of the M9GlcNAc1 structure (Figure 5; Supplementary Data 5). Complex *N*-glycans were most frequently represented by biantennary structures in melanocytes. Biantennary and tetraantennary *N*-glycans were similarly represented in melanocytes and primary and metastatic melanoma cells, accounting for 20.43%-24.55% and 0.18%-1.12%, respectively. Melanoma cells contained a statistically significant amount of triantennary structures, accounting for 29.45% and 24.88% of the *N*-glycome in primary and metastatic melanoma cells, respectively, compared to 5.77% in melanocytes (p < 0.05) (Figure 4, Supplementary Data 4).

**Figure 4.**
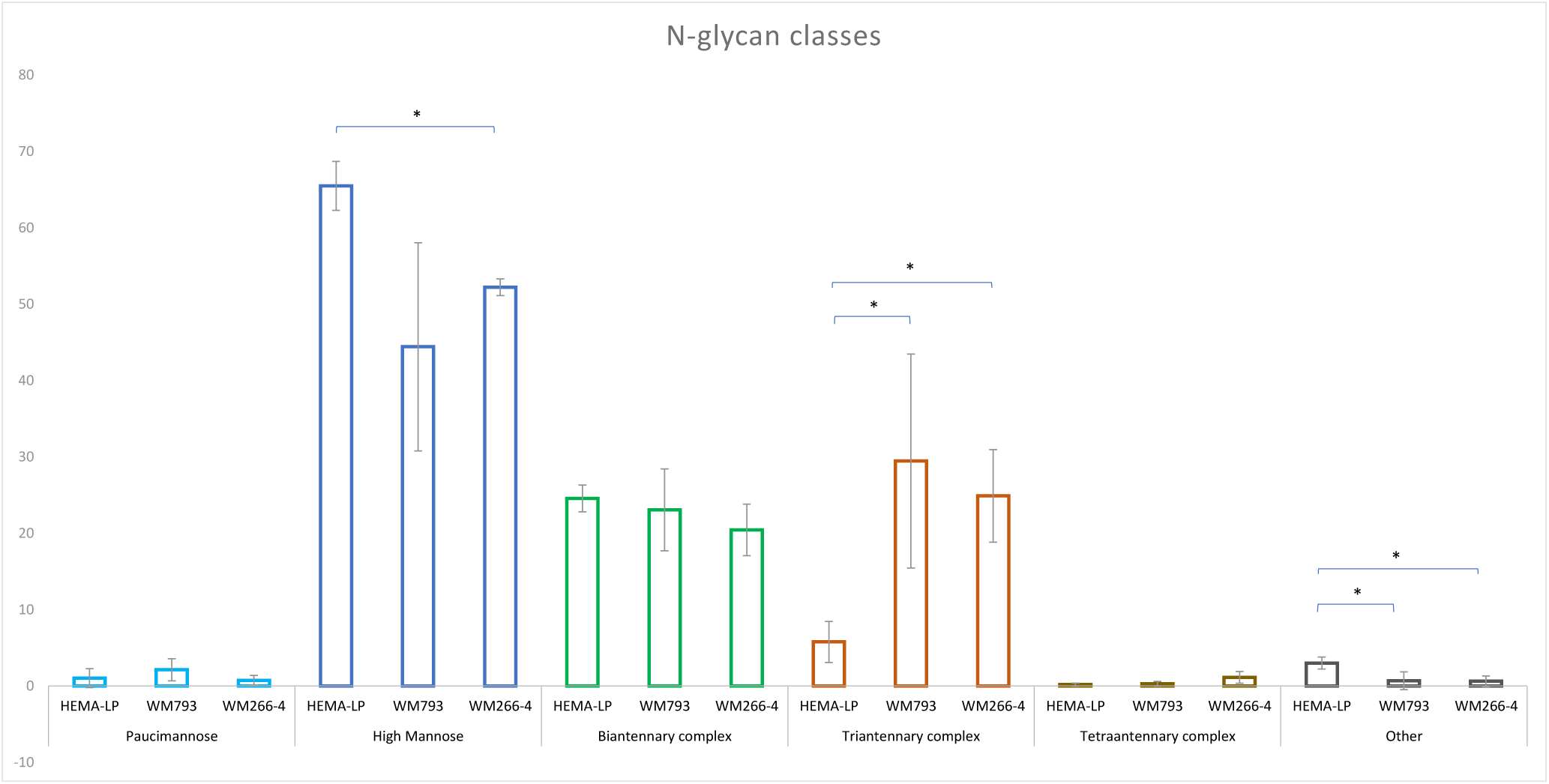
Relative percentage abundance of different classes of *N*-glycans based on MALDI-TOF MS analysis of EEA derivatized samples. The following glycan classes have been identified: paucimannose, high-mannose, diantennary complex, triantennary complex, and tetraantennary complex. The corresponding data are presented in Supplementary Data 4. Error bars represent standard deviation (n = 3). *p<0.05 (t-test in Excel).

**Figure 5.**
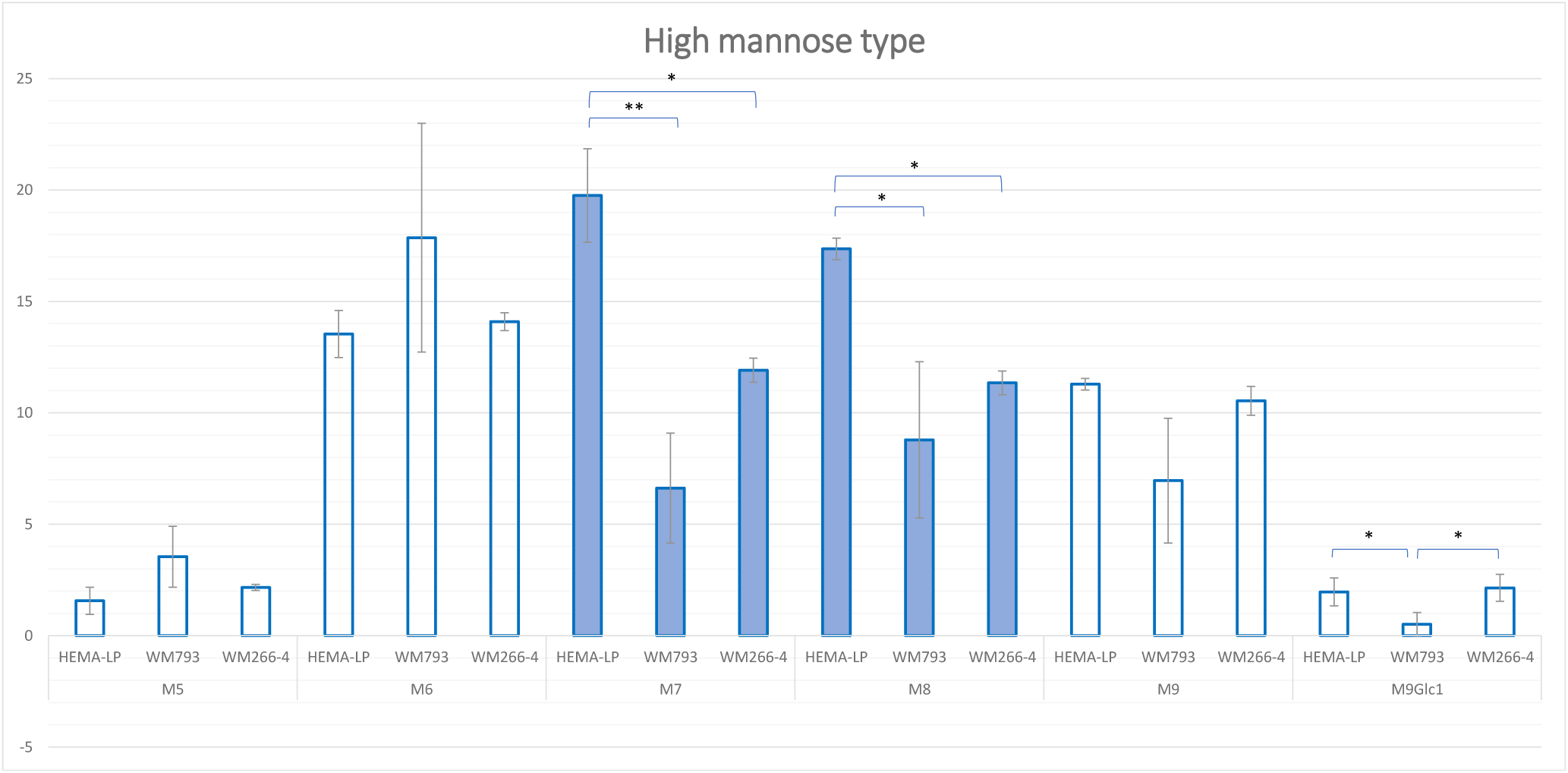
Relative percentage abundance of subclasses of high-mannose-type glycans (M5 – M9Glc1) based on MALDI-TOF MS of derivatized samples. The corresponding data are presented in Supplementary Data 5. Error bars represent standard deviation (n = 3). *p<0.05; **p<0.01(t-test in Excel).

### Other structures

Unexpected peaks remained on the chromatograms after digestion with the ABS+BTG+BKF+JBH+JBM and ABS+SPG+BKF+JBH+JBM enzyme arrays, which suggested the presence of Lewis epitopes. The fucose residue on the glycan structure’s antenna made it impossible to remove galactose with any galactosidase. To verify the presence of these epitopes, we conducted additional digestions using α1-3,4 fucosidases from *Ruminococcus gnavus* (RGF) or *Xanthomonas manihotis* (XMF), with or without BTG, SPG, or XMF. We also used α-galactosidase from green coffee beans (GCG) to check for the presence of α-linked galactose. These digestions were performed in a single run. The resulting chromatograms are shown in Supplementary Figures S5–S7. The obtained data are presented in Supplementary Table S8-S10.

HILIC HPLC analysis of products from the aforementioned digestions revealed Lewis X and A epitopes on *N*-glycans from all analyzed cell lines. Together with the results of previous digestions (see Supplementary Tables S1-S3), these findings suggest that Lewis epitopes are present in the F(6)A2BF(3)1G(4)1S(3)1, F(6)A3F(?)1G(4?,3)3-G(4)1S(6)2, and F(6)A4F(?)1G(4,4,?,3)4-G(4)1S(3)1 structures of melanocytes. These structures have been confirmed by other protocols. The first structure was confirmed by the presence of the 2408 m/z ion in the MALDI-TOF-MS analysis after ABS desialylation (see Supplementary Table S5). The other structures were confirmed by the presence of the 3162 m/z (GU 10.51 and 10.63) and 3236 m/z (GU 10.90) ions in the UPLC-MS analysis (see Supplementary Table S4), as well as the 3241 m/z ion in the MALDI-TOF-MS analysis after EEA derivatization (see Supplementary Table S6). Lewis epitopes were found on the F(6)A3F(3)1G1G(4)2-G(4)1S(6)2 and F(6)A3F(4)1G2G(4)1-G(4)1S(6)2 structures in WM793 melanoma cells, which was confirmed by the presence of a m/z 3241 signal in the MALDI-TOF-MS analysis after EEA derivatization (see Supplementary Table S6). However, these compounds were present in very low concentrations, as indicated by the weak signals in the EIC and mass spectra (see Supplementary Data 6). In the case of WM266-4 melanoma cells, Lewis epitopes were found on the F(6)A2[6]BF(3)1G(4)1 and F(6)A3F(4)1G2G(4)1-G(4)1S(6)2 compounds. The first structure was confirmed by the presence of the m/z 2408 ion in the MALDI-TOF-MS analysis after ABS desialylation (see Supplementary Table S5). The second structure was confirmed by the m/z 3241 signal in the MALDI-TOF-MS analysis after EEA derivatization (see Supplementary Table S6). Only a very low concentration was detected in the UPLC-MS analysis (see Supplementary Data 6).

Additional digestion with galactosidases, which hydrolyze β-glycosidic bonds, confirmed the presence of -G1-G1 motif in all analyzed cell lines. These unit was found on the F(6)A1[6]G1-G1S(6)2 compound, which was confirmed by the m/z 2364 ion in the MALDI-TOF-MS analysis after EEA derivatization (see Supplementary Table S6). For WM793 cells, this structure was likely also present in a monosialylated form, F(6)A1[6]G1-G1S(6)1 (*m/z* 2045; see Supplementary Table S6). For melanoma cells, an additional structure bearing the - G1-G1 unit was found (F(6)A3G(4)2G1-G1S2) and was confirmed by the m/z 3016 UPLC-MS analysis (see Supplementary Data 6). Interestingly, digestion with α-galactosidase from green coffee beans revealed the presence of α-galactose residue. Very low concentrations of it were seen only in the *N*-glycome of melanoma cells as the compound F(6)A3G(4)3Ga1S2 (*m/z* 3016; see Supplementary Data 3).

Furthermore, structures that contain Lewis epitopes (GU at 10.55, 10.63, and 10.88 for HEMA-LP, and GU at 10.58 and 10.62 for melanoma cells, as seen in Supplementary Tables 1-3, columns C-G) also carry the -GlcNAcβ1-4Galβ1-4Gal group (assigned G(4)1-G(4)1). This is confirmed by the disappearance of these peaks after digestion with a mixture of ABS+SPG+JBM or ABS+SPG+JBH+JBM (see Supplementary Tables 1–3, columns CP–CR, for HEMA-LP and WM266-4 or WM793 cells, respectively).

## Discussion

Each methodology we used enriched the glycoprofiling analysis, yielding largely consistent results. Observed differences mainly concern glycans present in smaller amounts. Glycoprofiling on TSKgel-Amide and BEH-Amide produced slightly different results (protocols 1 and 2) due to the polarity differences of the stationary phases and the additional purification steps before UPLC-MS analysis. Additionally, the internal structural features of different glycans may cause them to co-migrate under the same peak, and their relative abundance may vary depending on the sample repetition. However, the UPLC-MS technique allowed us to identify structures in which galactose is linked to another galactose residue. In turn, using MALDI-MS after EEA derivatization (protocol 4) increased the signal intensity and sensitivity by reducing the loss of labile terminal sialic acids and differentiating the position of the sialic acid linkage. Conversely, desialylating glycans before MALDI-MS analysis (protocol 3) reduces the size of glycans simplifying the profiles. Only this method allowed us to detect polylactosamine structures. Finally, by using arrays of several exoglycosidases and combining them with HILIC HPLC, we obtained detailed structural information about the glycans, including identification of the individual monosaccharides, their linkages, and their positions within the glycan structures. This methodology enabled the characterization of complex glycan heterogeneity and isomers that would be difficult to resolve by HILIC HPLC alone.

It is well known that melanoma *N*-glycomes dynamically changes during tumor progression toward a decrease in I-branched glycans and an increase in sialylation and fucosylation (Sweeney 2018, Agraval 2018, Niveau 2024). Abrahams demonstrated that high-mannose structures (M5-M9Glc1), primarily M8 and M9, made up 25.01% of the total *N*-glycan pool released from the melanoma tumor tissue of stage III lymph node patients and 24.94% of the total *N*-glycan pool released from the melanoma tumor tissue of stage IV lymph node patients. Among stage III melanoma patients, 18.81% of the *N*-glycans were bi-antennary, 17.67% were tri-antennary, and 8.49% were tetra-antennary. Among *N*-glycans isolated from stage IV melanoma patients, 16.90% were bi-antennary, 20.62% were tri-antennary, and 10.69% were tetra-antennary (Abrahams, 2016). Another report showed that high-mannose structures were the most abundant in WM266-4 (44.6 ± 3.1%) and WM115 (50.3 ± 2.5%) cells (Kinoshita et al., 2014). Our results are in agreement with the aforementioned studies and demonstrate that *N*-glycans released from melanocytes, as well as from primary and metastatic melanoma cells, are most frequently represented by oligomannose structures (M5-M9), accounting for approximately 65%, 44%, and 52% of detected *N*-glycans, respectively. However, normal cells were characterized by a statistically significantly higher expression of M7 and M8 mannose-type glycans compared to melanoma cells. The levels of bi- and tetra-antennary complex-type glycans were similar in all analyzed cell lines (approximately 23% and 0.2– 1.1%, respectively). There was a tendency for their concentrations to decrease and increase, respectively, in metastatic cells. The greatest differences in expression were observed for triantennary complex-type structures, favoring melanoma cells (p < 0.05).

We demonstrated that melanoma cell *N*-glycans are more sialylated than melanocytes. Melanoma cells contained a large proportion of *N*-glycans with α2-6-linked sialic acid, while melanocytes mainly had α2-3-linked sialic acid in their released oligosaccharides. These findings align with those of Abrahams, who observed a higher overall abundance of α2-6-linked sialic acid residues compared to α2-3-linked residues in *N*-glycans released from the total membrane proteins of the metastatic melanoma cell line MM253. The same study, which analyzed *N*-glycans released from lymph node melanoma tumor tissue of stage III/IV patients, showed that the poor prognostic group had a higher amount of α2-6-linked mono- and disialylated residues than the good prognostic group, which had an overall higher abundance of α2-3-linked sialic acid residues (Abrahams, 2016). However, analyses of sialylation in selected adhesion proteins yielded contradictory results. For example, loss of α2-3-linked sialic acid in the cell adhesion molecule L1CAM was observed during the transition of melanoma cells from VGP to a metastatic stage (Hoja-Łukowicz et al., 2013). Another study observed a reduction in α2-6-linked sialic acid expression and an increase in α2-3 sialylation of both αvβ3 integrin subunits in melanoma progression (Pocheć et al., 2015).

Agraval demonstrated that core fucosylation is significantly increased in metastatic melanoma and can promote metastasis by influencing proteins such as L1CAM (Agraval et al., 2017). Abrahams reported that complex-type glycans with core fucosylation comprised 27-55% of the glycan population in melanoma. Our results showed that approximately 20% of the total oligosaccharide pool is core fucosylated in melanoma cells. Melanocytes did not differ from this level, with about 24% core fucosylation (see Supplementary Figure S8 and Supplementary Data 7).

We demonstrated the presence of the sialyl Lewis X antigen in melanocytes and metastatic melanoma cells. Previous studies have shown that C57BL/6 mice that were injected with melanoma cells that expressed moderate amounts of sialyl Lewis X in poly-N-acetyllactosamine (B16-FTIII·M cells) developed large numbers of lung tumor nodules. Conversely, C57BL/6 mice that were injected with melanoma cells that expressed the highest levels of sialyl Lewis X in shorter *N*-glycans (B16-FTIII·H cells) developed few lung nodules. The same researchers obtained similar results while working with the MeWo human malignant melanoma cell line. Cells expressing high levels of sialyl Lewis X were not metastatic; however, those corresponding to B16-FTIII·M cells were highly metastatic. It has been suggested that the formation of metastases depends on the acquisition of the sialyl-Lewis X antigen and the amount and structure of the backbone glycans containing sialyl Lewis X (Ohyama et al., 1999). We also detected complex-type glycans carrying the sialyl-Lewis A antigen. This finding aligns with a previous study suggesting that sialyl-Lewis A is a progression marker in melanocytic lesions because its expression increases with the thickness of primary melanoma lesions and is expressed in approximately 90% of metastatic lesions.

Previous studies have observed changes in the expression of the LacdiNAc group during oncogenic transformation. Enhanced expression of the LacdiNAc group appears to be responsible for various effects depending on tumor type (Hirano et al., 2014). Hsu et al. (2011) presented evidence that LacdiNAc group expression inhibits human neuroblastoma cell migration and invasion. However, the reverse relationship was found in human prostate, ovarian, and pancreatic cancers, where enhanced LacdiNAc expression has been associated with tumor progression (Hirano et al., 2014). The LacdiNAc motif has previously been identified on bi-antennary glycans in Bowes melanoma tissue plasminogen activator (Chan et al., 1991). Abrahams suggested that the LacdiNAc motif present on the MCAM glycoprotein could serve as a potential melanoma diagnostic marker (Abrahams, 2016). Our results confirmed the absence of the LacdiNAc glycoepitope in melanocytes and its presence in melanoma cells. This supports the idea that this glycoepitope can distinguish normal cells from melanoma cells.

In this study, we compared the chromatograms of HEMA-LP, WM793, and WM266-4 oligosaccharides that were processed with sequential exoglycosidase digestion using galactosidases BTG and SPG. We found that complex *N*-glycans containing β1-3-linked galactose are characteristic of melanoma cells, while such structures are rare among melanocyte-derived oligosaccharides. Furthermore, we observed a statistically significant increase in the presence of type 1 LacNAc in metastatic cells compared to primary melanoma cells. A previous study found the presence of β1-3-linked galactose on L1CAM *N*-glycans isolated from metastatic melanoma cells (WM1205Lu cell line). This study showed that one-fourth of all structures possessed a Galβ1-3-linked residue to a GlcNAc residue (type 1 LacNAc). Furthermore, the ratio of type 1 LacNAc to type 2 LacNAc was higher in the metastatic L1CAM glycan pool (WM1205Lu cell line) than in the primary L1CAM glycan pool (WM793 cell line) for digalactosylated, biantennary complex-type oligosaccharides. This study also found that β1-3-linked galactose residues were present almost exclusively on the 6-arm of complex-type glycans. Therefore, the presence of a site-specific β3-galactosyltransferase in melanoma has been suggested (Hoja-Łukowicz et al., 2013). This study also demonstrated that β1-3-linked galactose is mainly present on the 6-arm in core structures. Taken together, these findings suggest that type 1 LacNAc may be a potential diagnostic marker of melanoma progression.

In a previous study, we reported the presence of the Galβ1-4Galβ1-4GlcNAc-motif on sialylated monoantennary complex-type glycans in L1CAM from melanoma cells at high amounts (about 16%) (Hoja-Łukowicz, 2013). The study identified this motif as a melanoma-associated carbohydrate antigen that is absent from normal cell surfaces. Interestingly, we detected small amounts of this structural motif on sialylated, multi-antennary, complex-type glycans. Our results do not contradict previous findings because they concern a different carrier structure. Furthermore, it is known that differences in the glycosylation patterns of normal and cancerous cells are primarily quantitative rather than qualitative. We also identified trace amounts of the Galβ1-3Galβ1-3GlcNAc- and Galα1-?Galβ1-4GlcNAc-motifs in all and melanoma samples, respectively. To the best of our knowledge, the former was detected for the first time. The latter was the result of bovine serum α1,3-galactosyltransferase (α3GalT) activity; calf serum was used to supplement the cell culture medium.

In conclusion, we established that type 1 LacNAc and LacdiNAc are promising structural motifs that can serve as glycomarkers for the early stages of melanoma progression. Further research is needed to more precisely determine the levels of these epitopes at different stages of melanoma progression.

## Conflict of interests

None of the authors has any competing interests in the manuscript.

## Acknowledgements

The authors thank Carolien Koeleman and Agnes Hipgrave Ederveen for excellent analytical support. This work was supported by the National Science Centre, Poland (grant number 2016/21/B/NZ3/00348).

## Author Contributions

K.G: Formal analysis, Writing – review and editing. P.L.-L: Data curation, Formal analysis, Methodology – review and editing. A.A: Data curation, Formal analysis, Methodology – review and editing. A.C: Data curation, Formal analysis, Methodology – review and editing. M.W: Data curation, Formal analysis, Methodology – review and editing. D.H-Ł: Conceptualisation, Data curation, Formal analysis, Methodology, Acquired the financial support– review and editing.

---

Supplementary Figure S1. Example sets of HILIC HPLC chromatograms of the SPS glycosidase series for melanocyte and melanoma cell samples. The corresponding data are presented in Supplementary Tables S1–S3 (columns B–G, P–U, and BH–CC). The used enzymes are ABS (α2-3,6,8 sialidase), SPG (β1-4 galactosidase), BKF (α1-6 fucosidase), JBH (β1–2,3,4,6 hexosaminidase), JBM (α1–2,3,6 mannosidase), and HPM (β1–4 mannosidase). Dotted lines indicate shifts in glycans digested by subsequent enzyme arrays. The structure abbreviations are the same as those in Table 2.

Supplementary Figure S2. UPLC-MS chromatograms for N-glycans released from melanocytes. The glycans detected for each peak are presented in Supplementary Table S4.

Supplementary Figure S3. UPLC-MS chromatograms for N-glycans released from primary WM793 melanoma cells. The glycans detected for each peak are presented in Supplementary Table S4.

Supplementary Figure S4. UPLC-MS chromatograms for N-glycans released from metastatic WM266-4 melanoma cells. The glycans detected for each peak are presented in Supplementary Table S4.

Supplementary Figure S5. Detection of Lewis X/A epitopes and the Galβ1-3Galβ1-3GlcNAc-motif (G1-G1) in *N*-glycans released from melanocytes using HILIC HPLC. The enzymes used are ABS (α2-3,6,8 sialidase), SPG (β1-4 galactosidase), BKF (α1-6 fucosidase), JBH (β1–2,3,4,6 hexosaminidase), JBM (α1–2,3,6 mannosidase), BTG (β1-3/4 galactosidase), RGF (α1–3,4 fucosidase), XMG (β1–3 galactosidase), and GCG (α-galactosidase). The corresponding data are presented in Supplementary Table S8. Dotted lines indicate the shifts of the glycans digested by the subsequent enzyme array.

Supplementary Figure S6. Detection of Lewis X/A epitopes, the Galβ1-3Galβ1-3GlcNAc-motif (G1-G1), and α-linked galactose in *N*-glycans released from primary WM793 melanoma cells using HILIC HPLC. The enzymes used are ABS (α2-3,6,8 sialidase), SPG (β1-4 galactosidase), BKF (α1-6 fucosidase), JBH (β1–2,3,4,6 hexosaminidase), JBM (α1– 2,3,6 mannosidase), BTG (β1-3/4 galactosidase), RGF (α1–3,4 fucosidase), XMG (β1–3 galactosidase), XSF (α1–3,4 fucosidase), and GCG (α-galactosidase). The corresponding data are presented in Supplementary Table S9. Dotted lines indicate the shifts of the glycans digested by the subsequent enzyme array.

Supplementary Figure S7. Detection of Lewis X/A epitopes and the Galβ1-3Galβ1-3GlcNAc-motif (G1-G1), and α-linked galactose in *N*-glycans released from metastatic WM266-4 melanoma cells using HILIC HPLC. The enzymes used are ABS (α2-3,6,8 sialidase), SPG (β1-4 galactosidase), BKF (α1-6 fucosidase), JBH (β1–2,3,4,6 hexosaminidase), JBM (α1– 2,3,6 mannosidase), BTG (β1-3/4 galactosidase), RGF (α1–3,4 fucosidase), XMG (β1–3 galactosidase), and GCG (α-galactosidase). The corresponding data are presented in Supplementary Table S10. Dotted lines indicate the shifts of the glycans digested by the subsequent enzyme array.

Supplementary Figure S8. Relative percentage abundance of fucosylated structures detected by MALDI-TOF MS after EEA derivatization of sialic acids. The corresponding data are presented in Supplementary Data 7.

## Notes

### Competing Interest Statement

The authors have declared no competing interest.

