## Supplementary material for "Type 1 LacNAc and LacdiNAc on *N*-glycans as molecular biomarkers of human melanoma cells": Supplementary Data 6.pdf

**HEMA-LP(3)**

H7N5F2S2-2AB (e.g. F2A3G(?)3G(?)1S2-2AB)

theoretical m/z $[M+2H]^{2+}$ =1582.0836

FLR

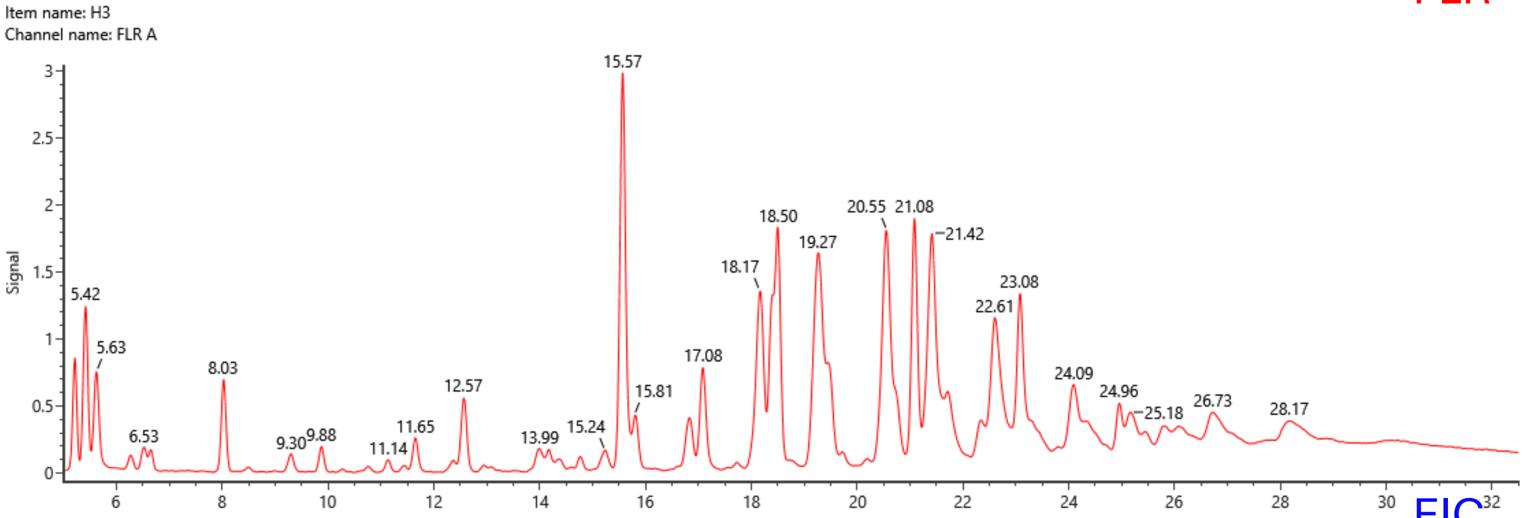


EIC

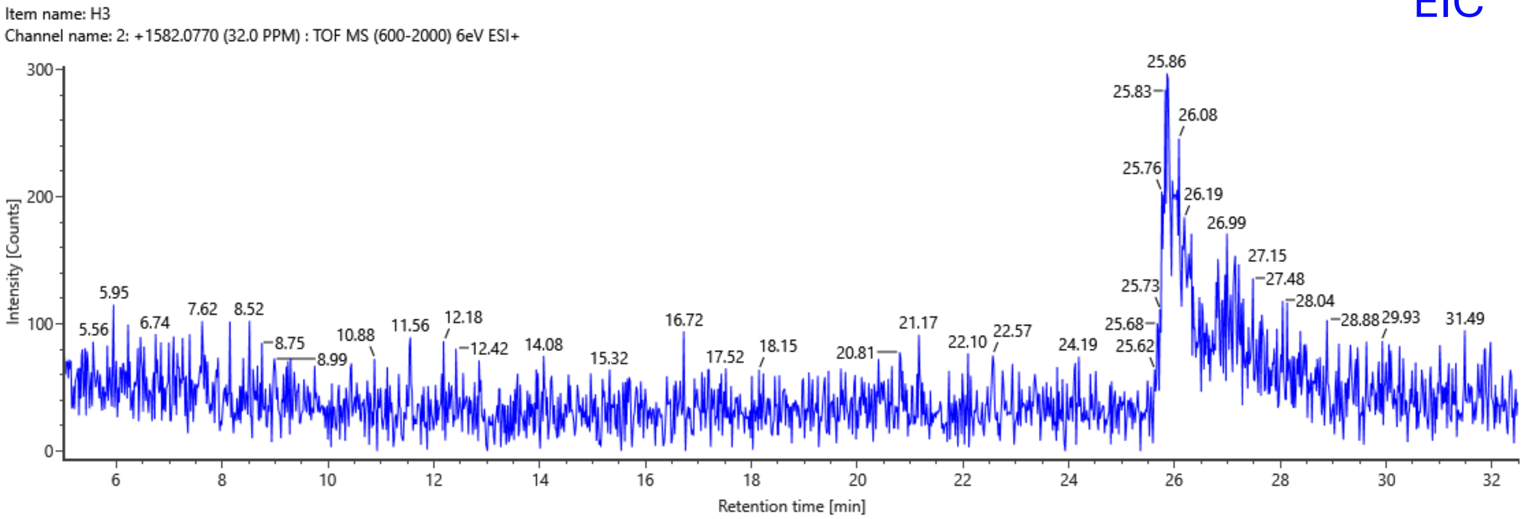


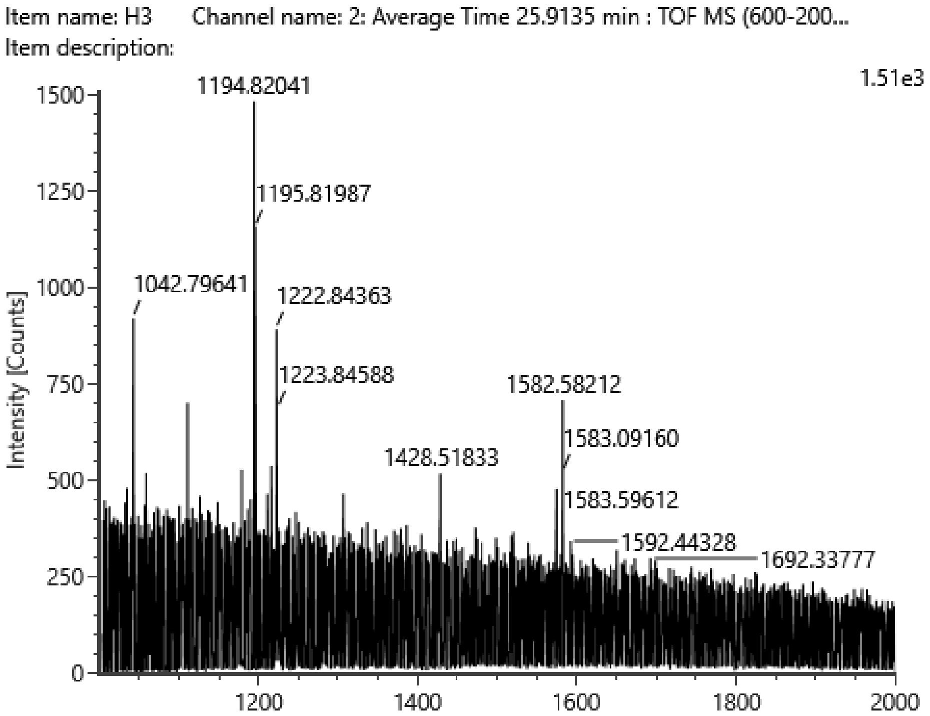


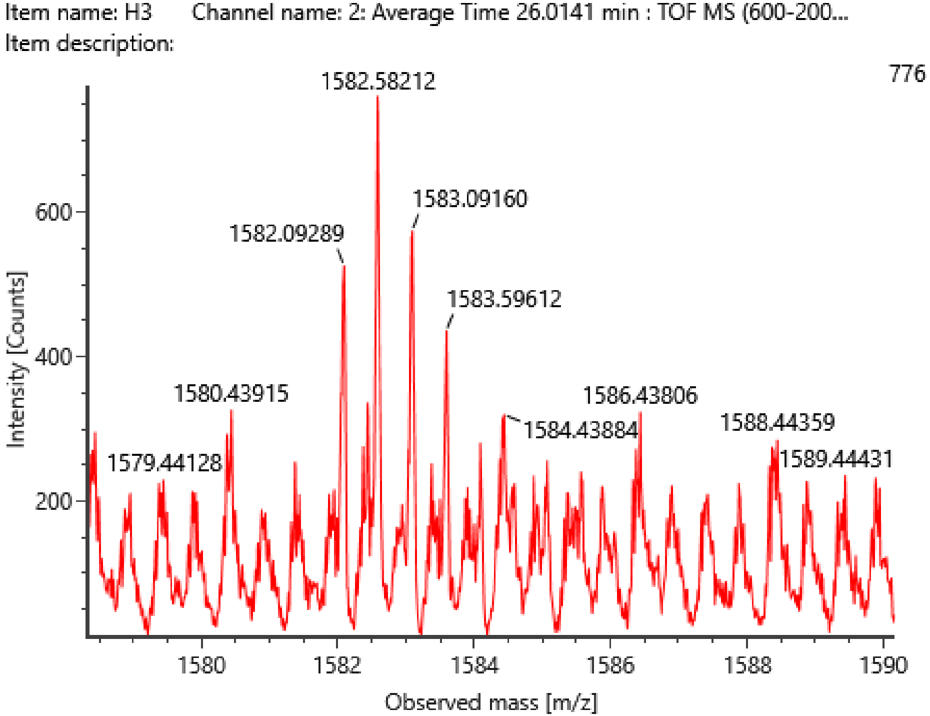

### HEMA-LP(3)

H8N6F2S1-2AB (e.g. F2A4G(?)4G(?)1S1-2AB)
theoretical m/z $[M+2H]^{2+}$ =1619.1020

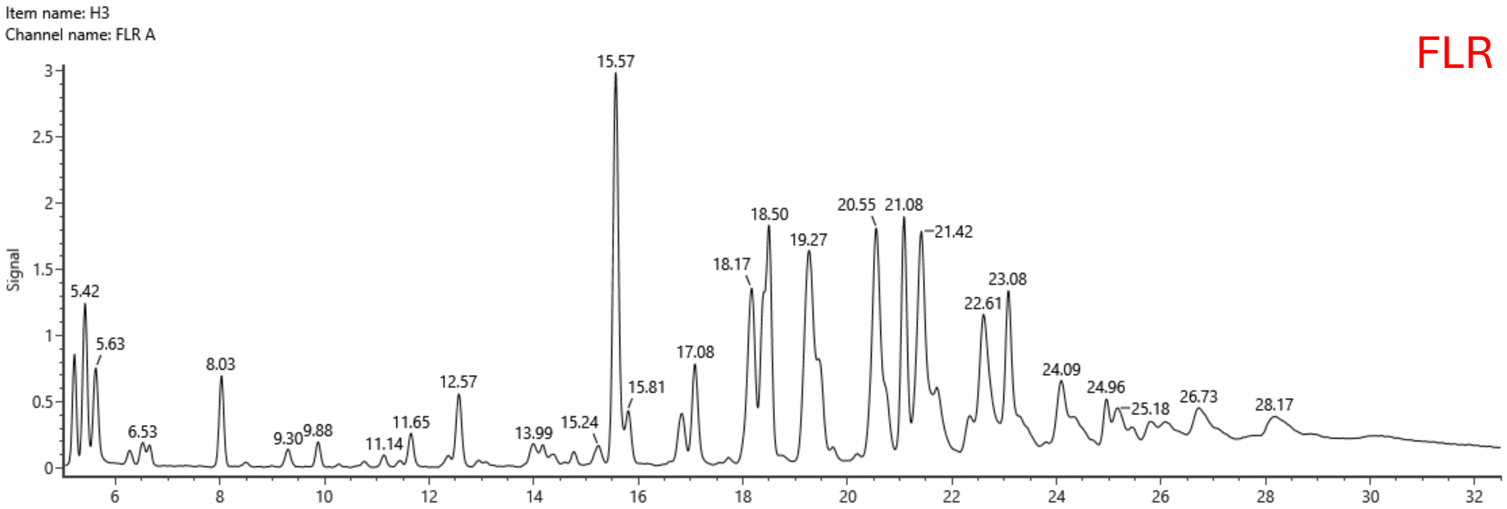


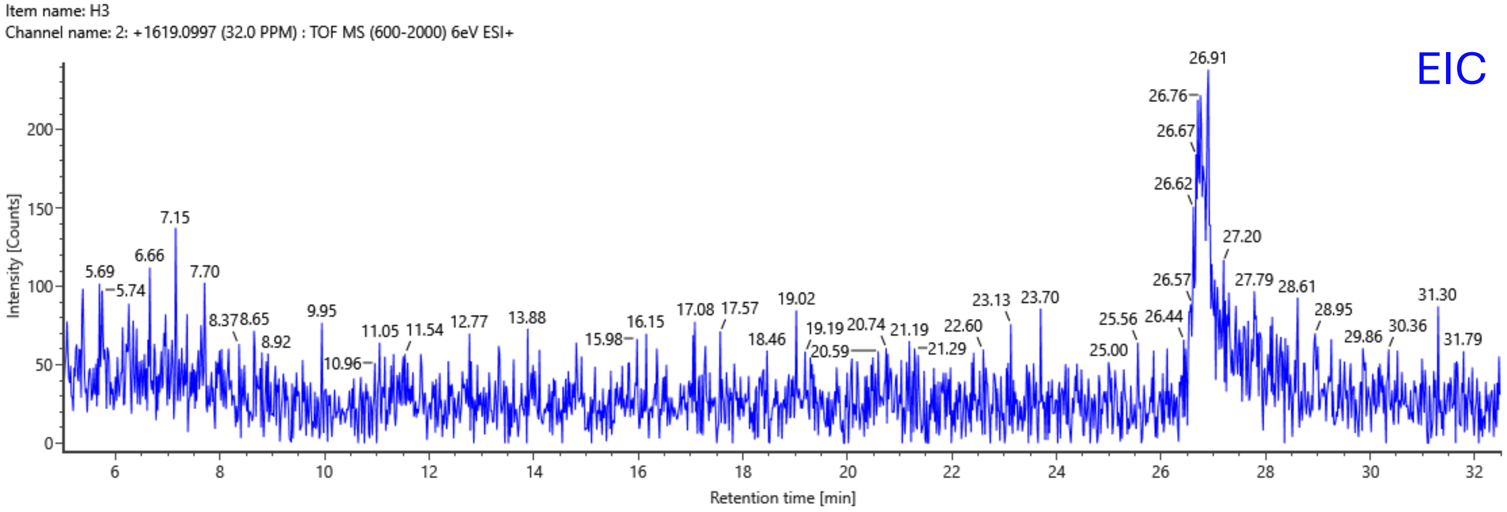


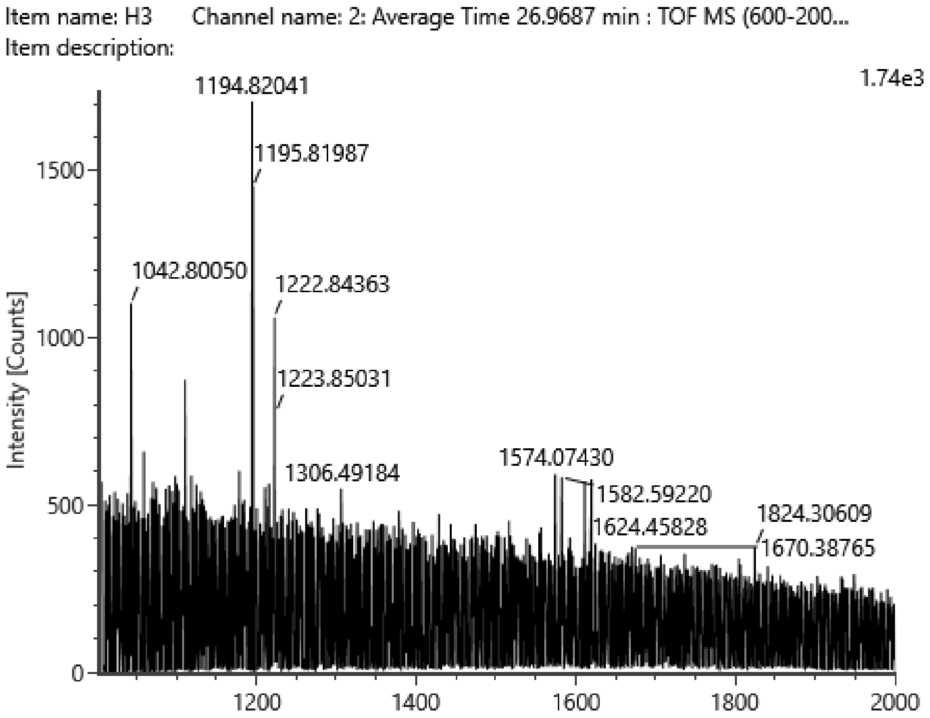


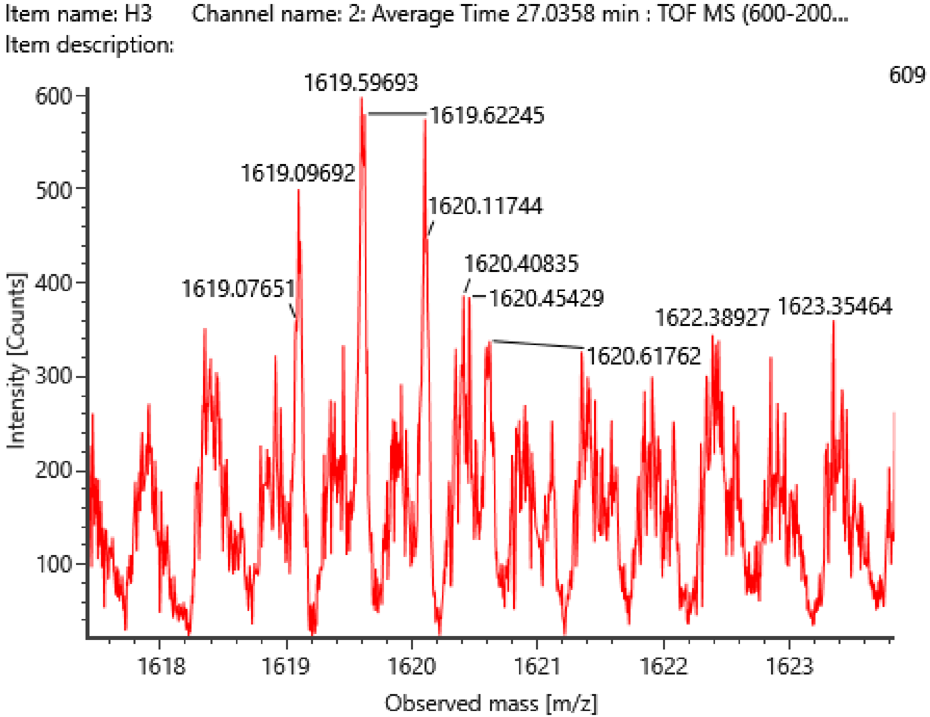

**WM793(1)**

H7N5F1S2-2AB (e.g. F1A3G(?)3G(?)1S2-2AB)

theoretical m/z $[M+2H]^{2+}$ =1509.0546

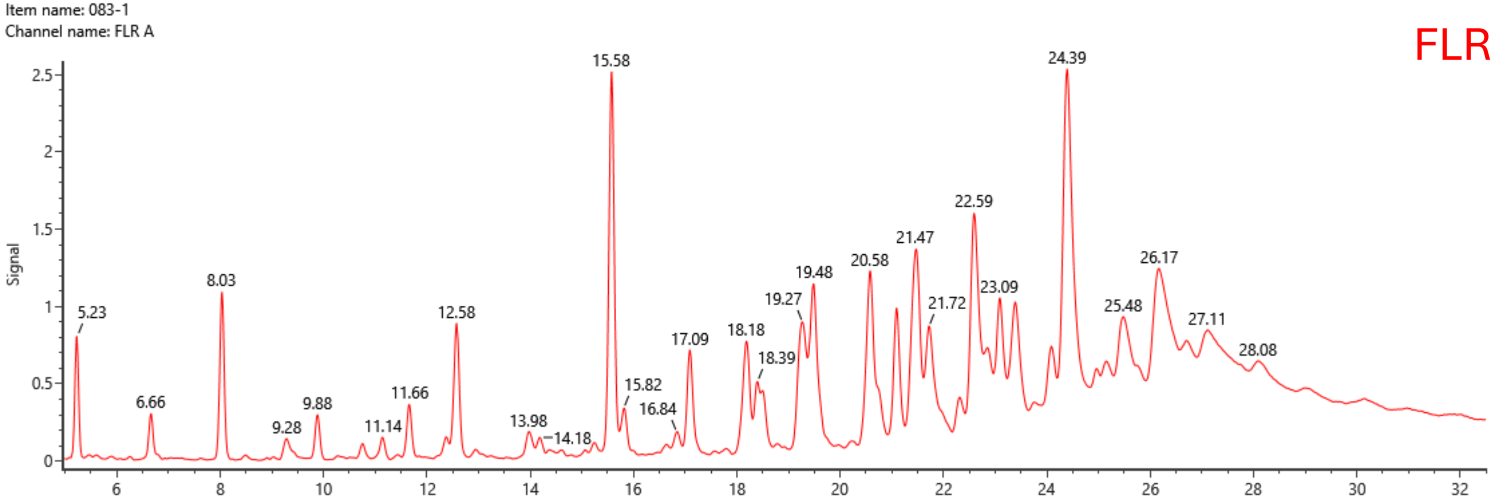


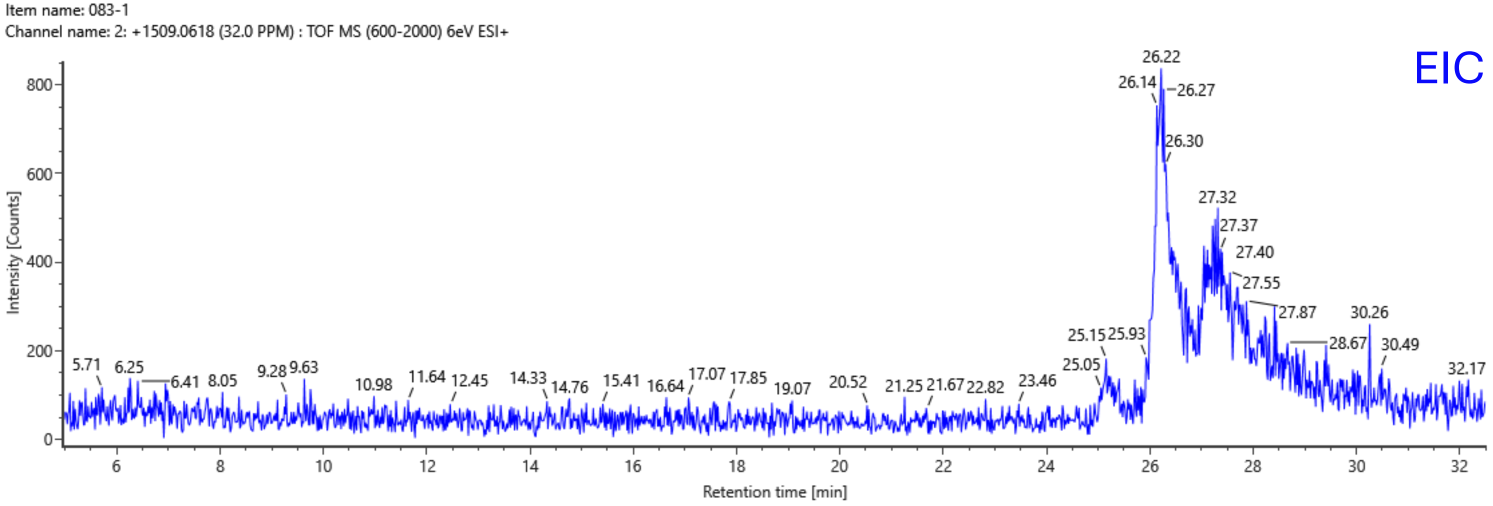


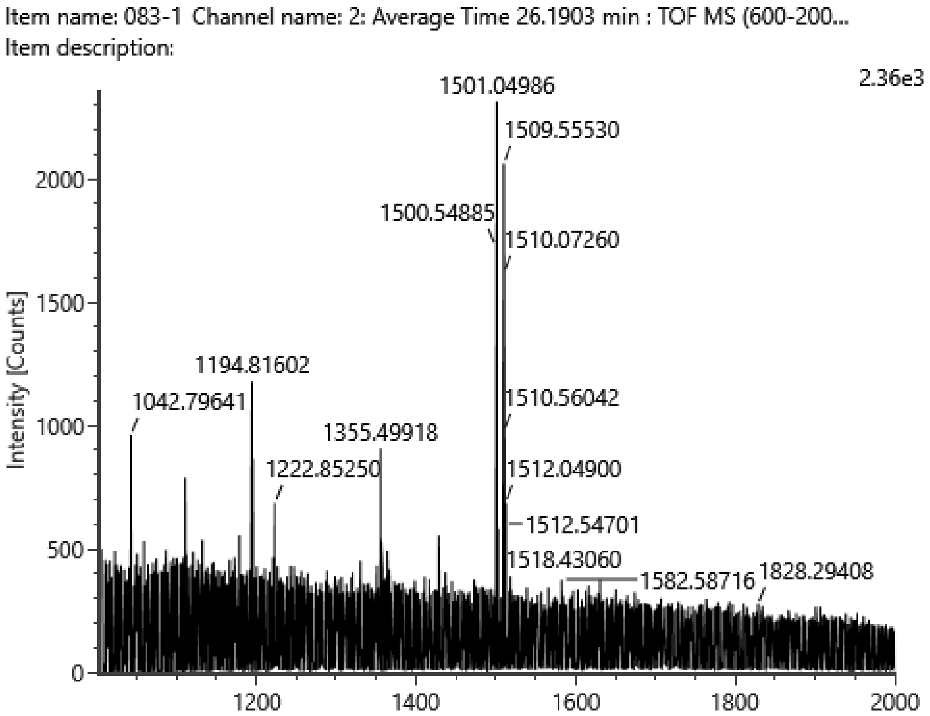


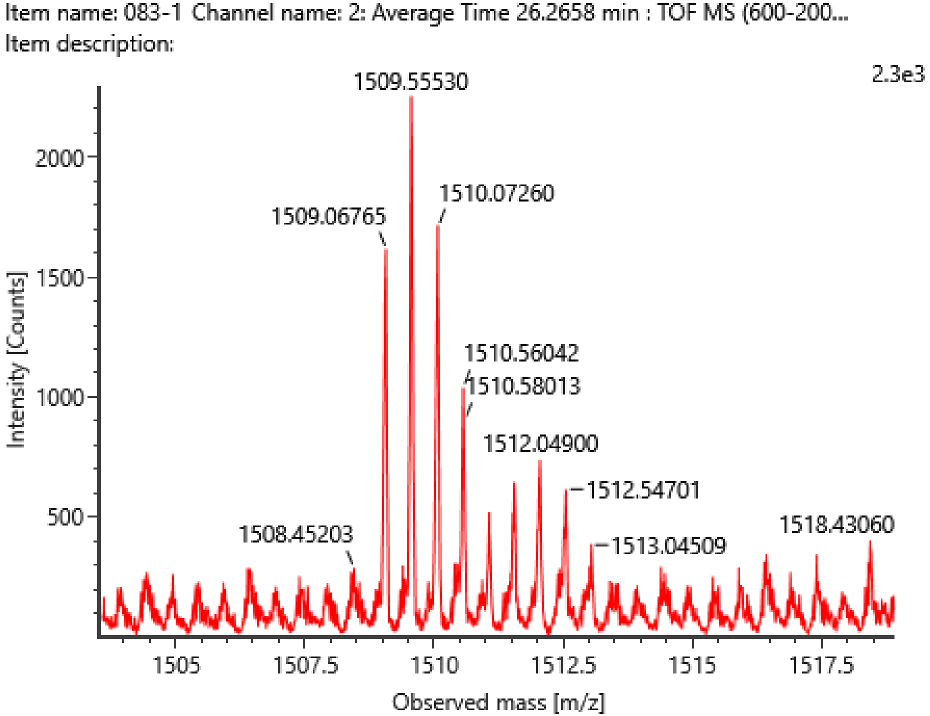

# WM793(1)

H7N5F2S2-2AB (e.g. F2A3G(?)3G(?)1S2-2AB)

theoretical m/z $[M+2H]^{2+}$ =1582.0836

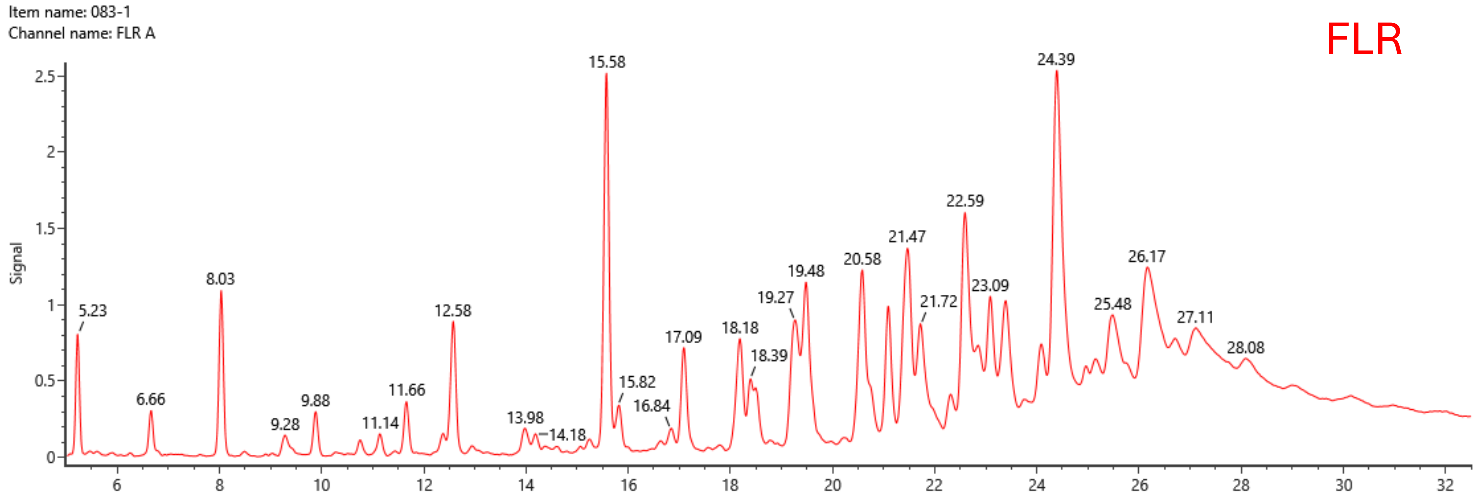


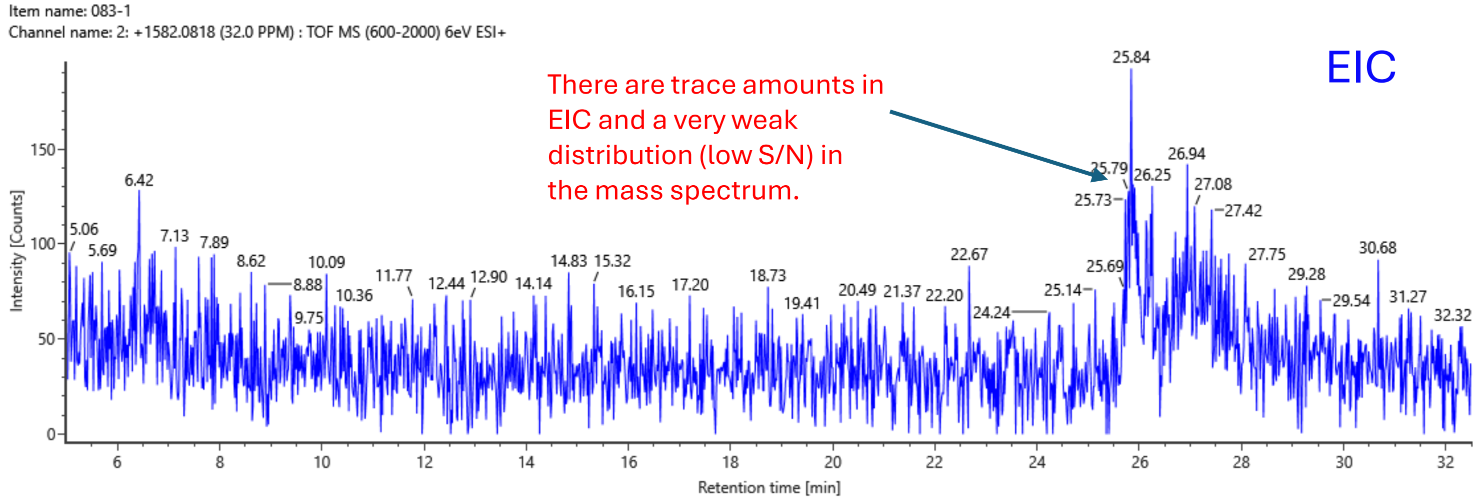


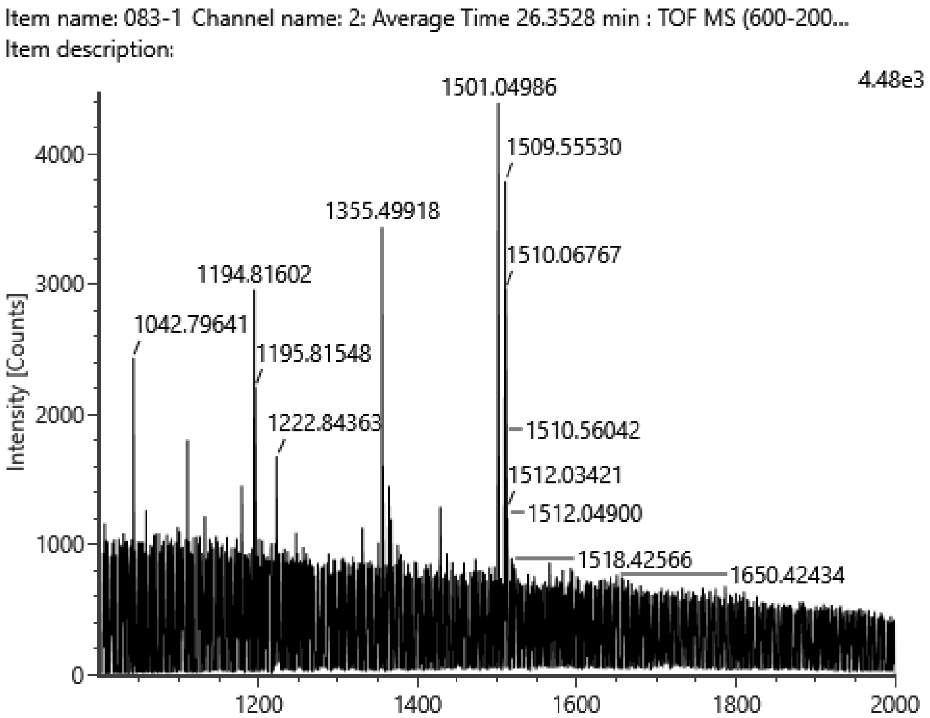


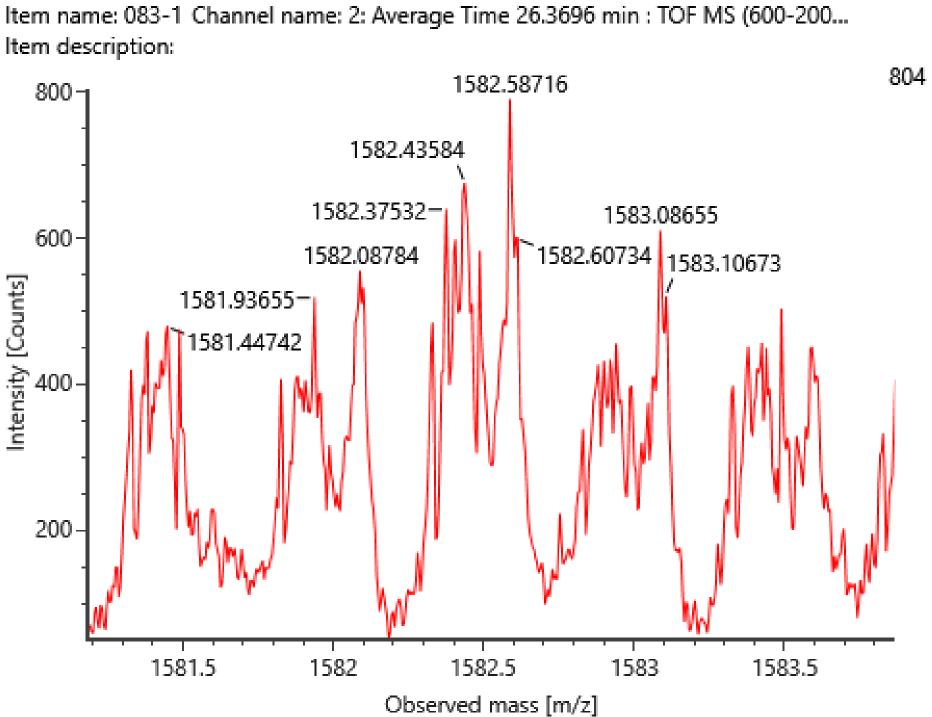

**WM793(1)**

H8N6F2S1-2AB (e.g. F2A4G(?)4G(?)1S1-2AB)

theoretical m/z $[M+2H]^{2+}$ =1619.1020

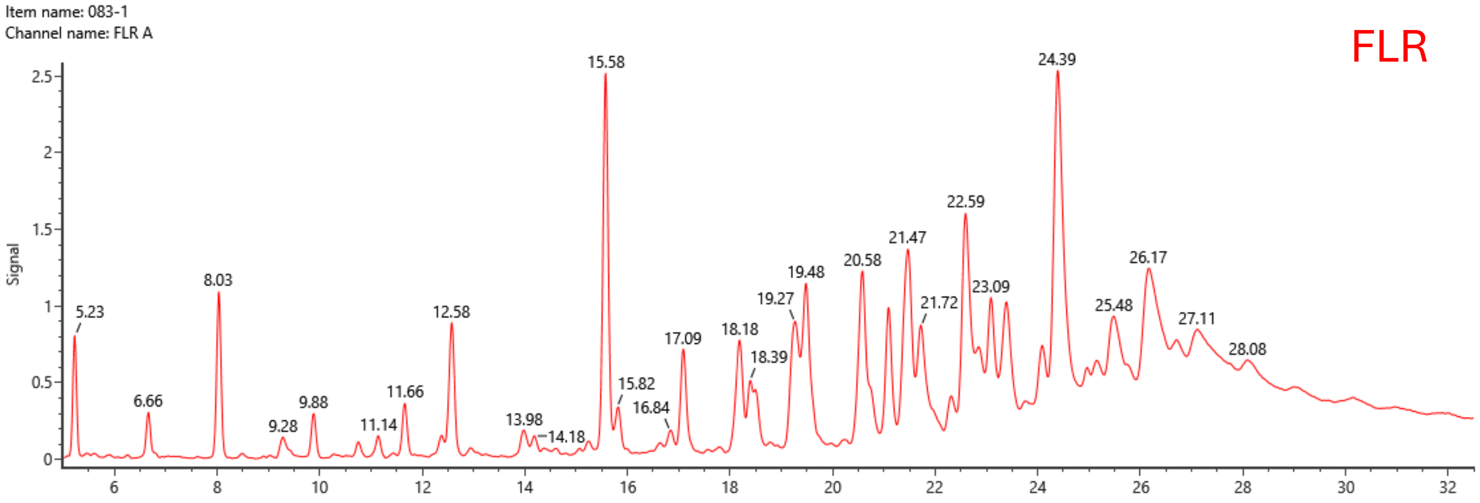


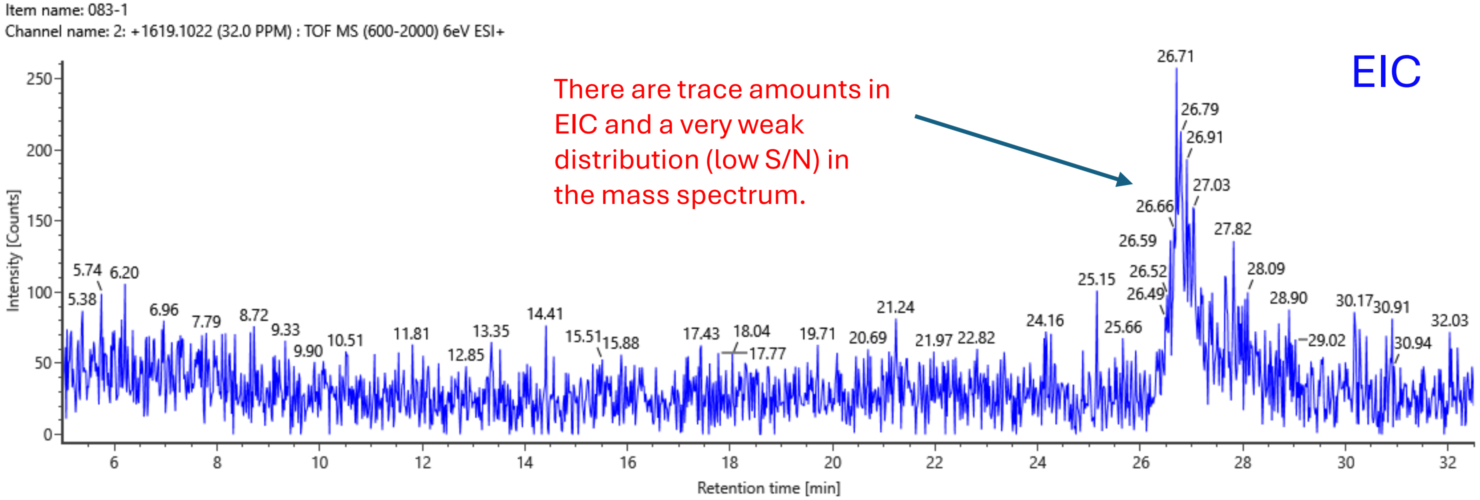


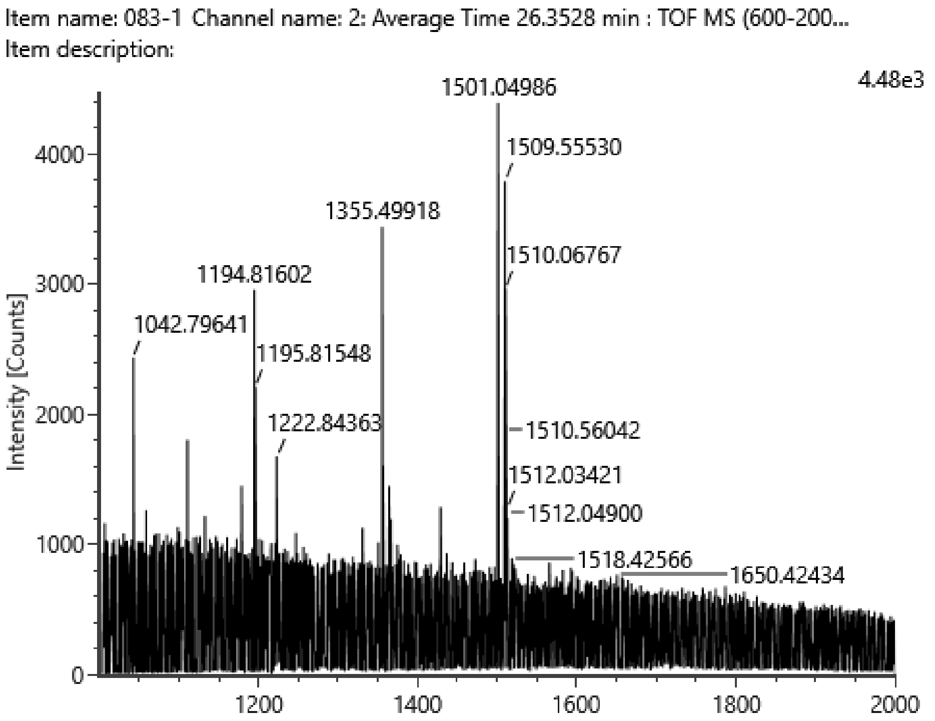


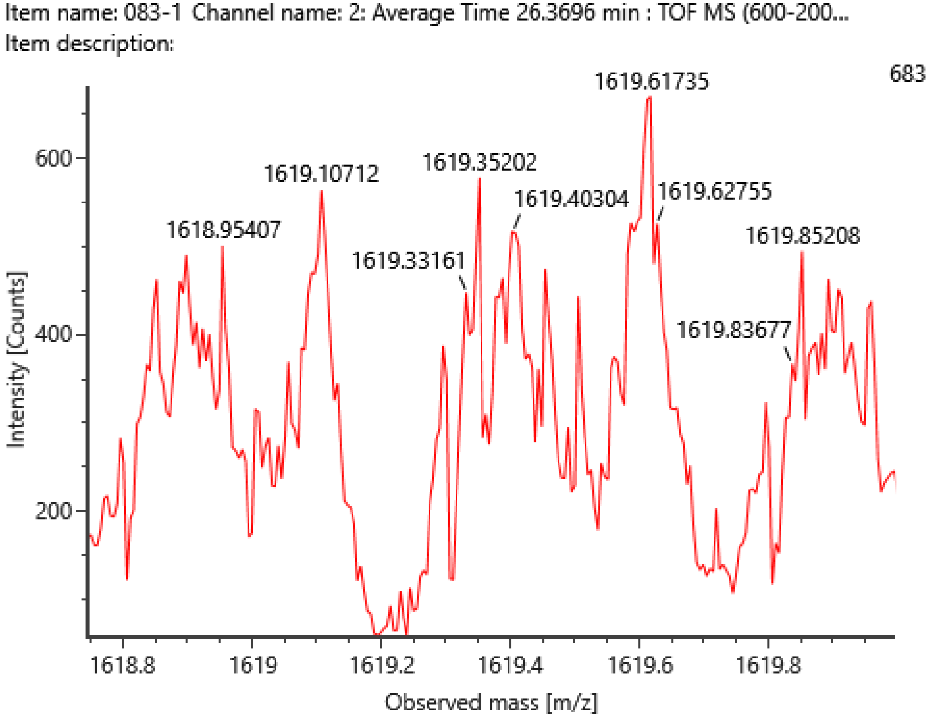

## WM266-4(1)

H7N5F1S2-2AB (e.g. F1A3G(?)3G(?)1S2-2AB)
theoretical m/z $[M+2H]^{2+}$ =1509.0546

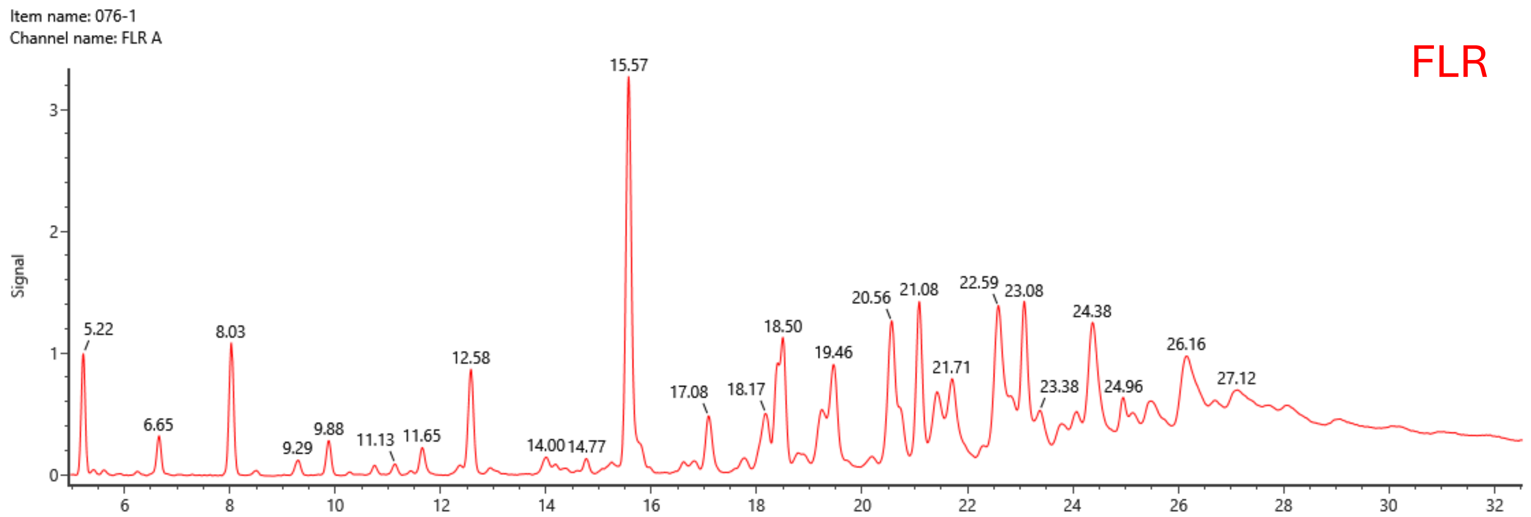


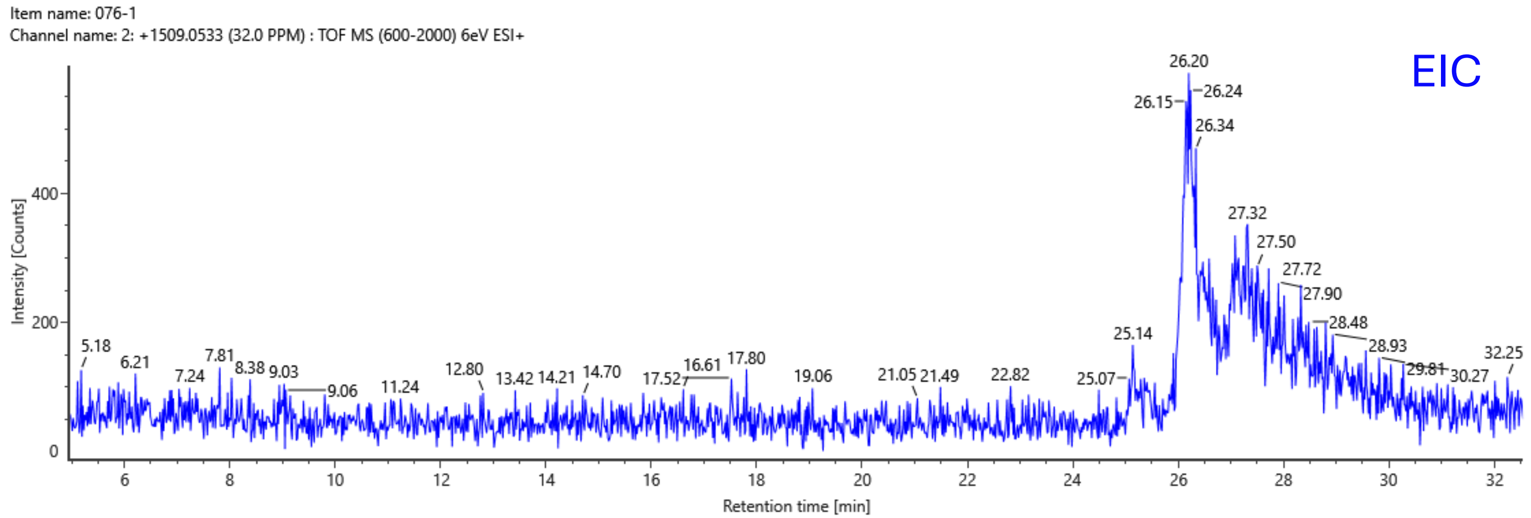


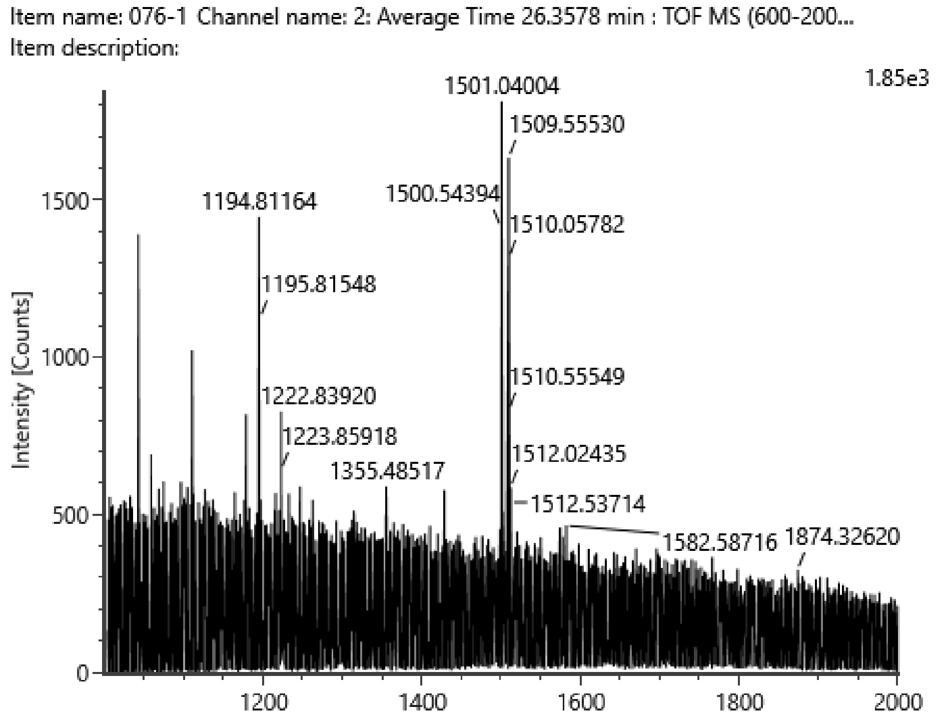


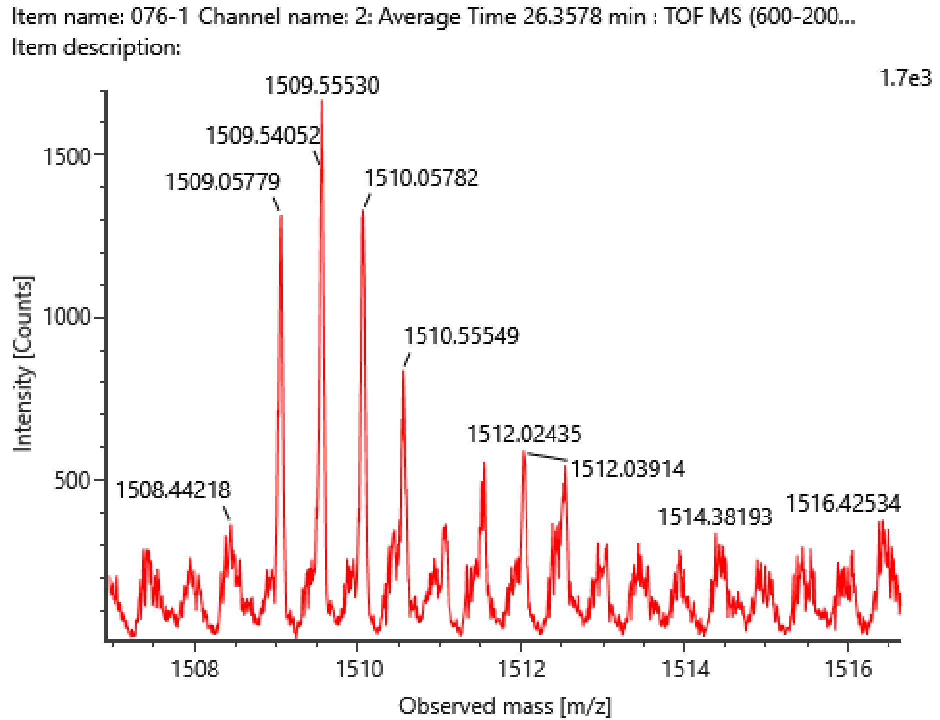

# WM266-4(1)

H7N5F2S2-2AB (e.g. F2A3G(?)3G(?)1S2-2AB)
theoretical m/z $[M+2H]^{2+}$ =1582.0836

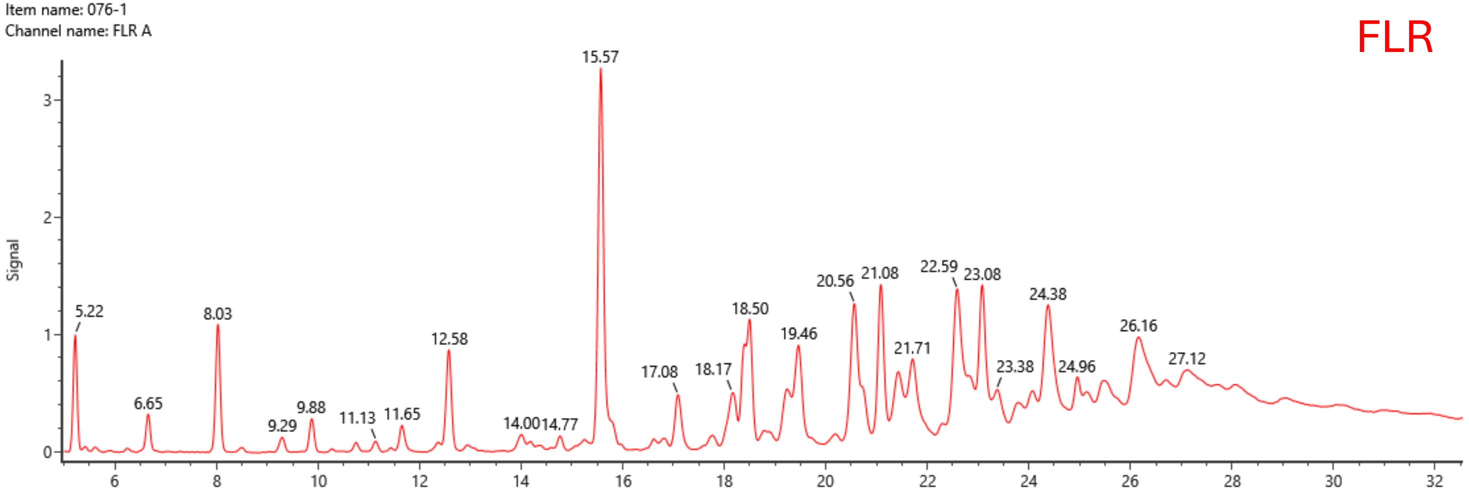


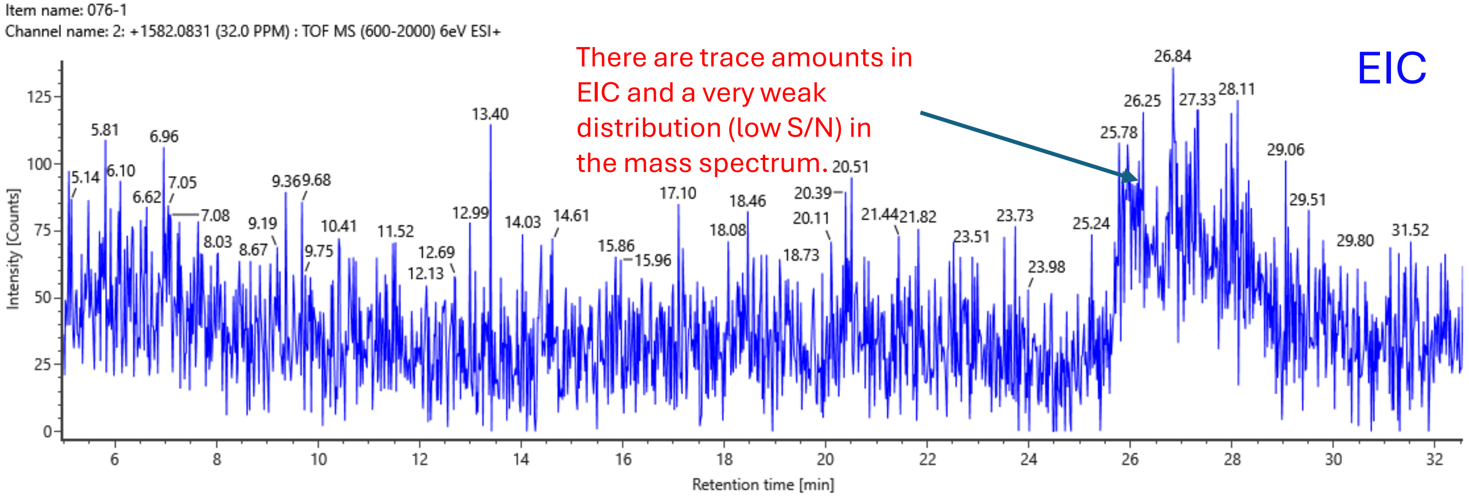


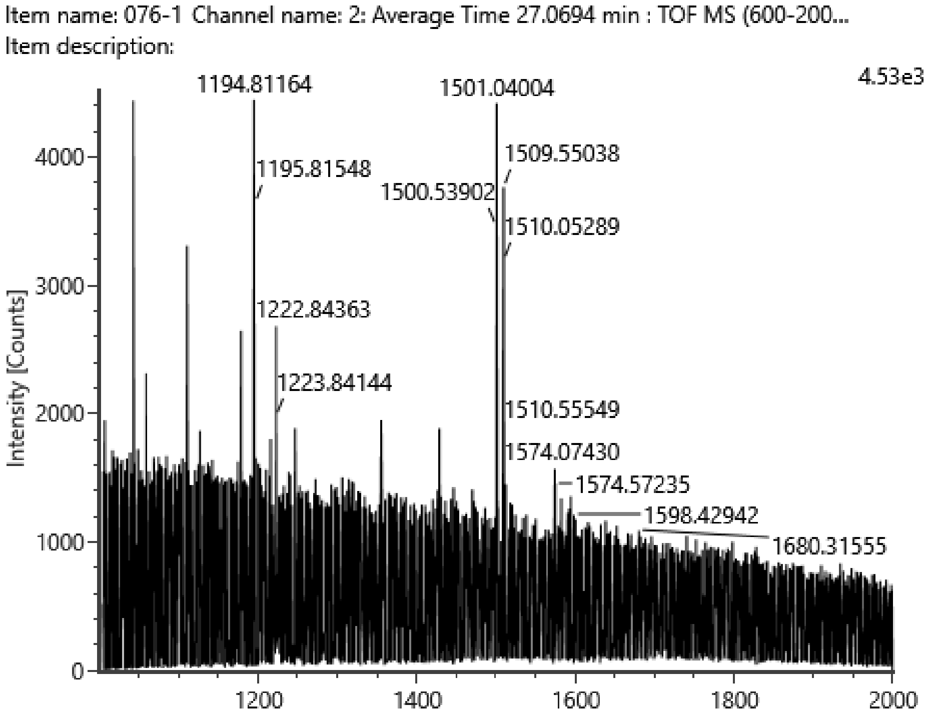


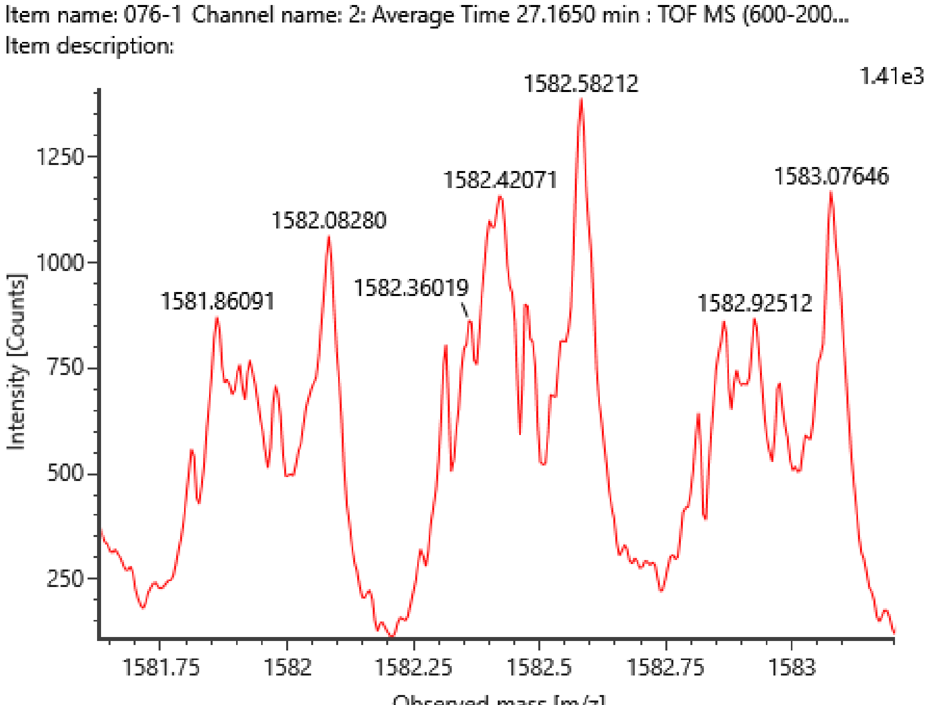

# WM266-4(1)

H8N6F2S1-2AB (e.g. F2A4G(?)4G(?)1S1-2AB)
theoretical m/z $[M+2H]^{2+}$ =1619.1020

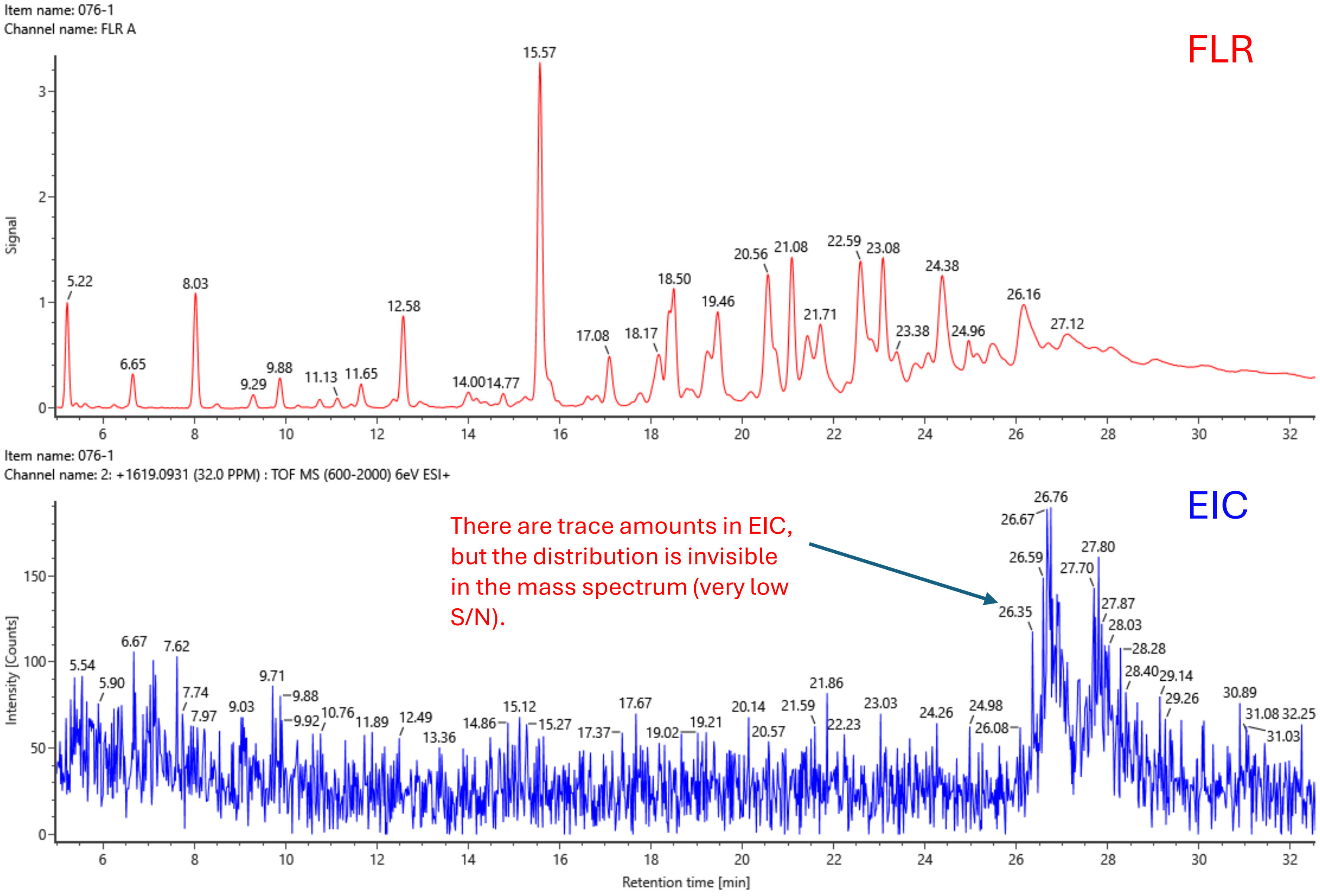


There are trace amounts in EIC, but the distribution is invisible in the mass spectrum (very low S/N).

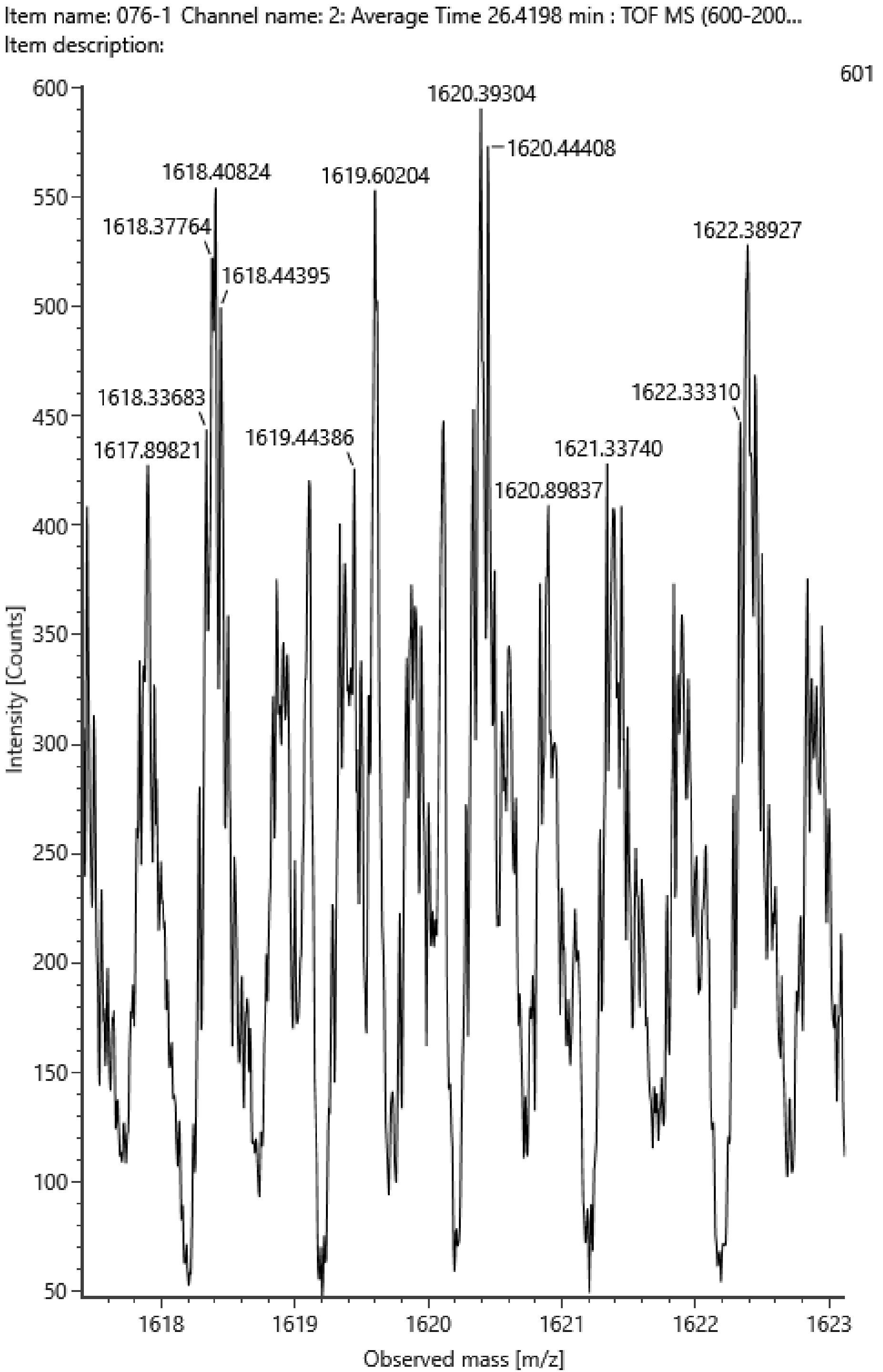
