## Supplementary figures and images for "Type 1 LacNAc and LacdiNAc on *N*-glycans as molecular biomarkers of human melanoma cells"

### Supplementary Figures S1.pdf

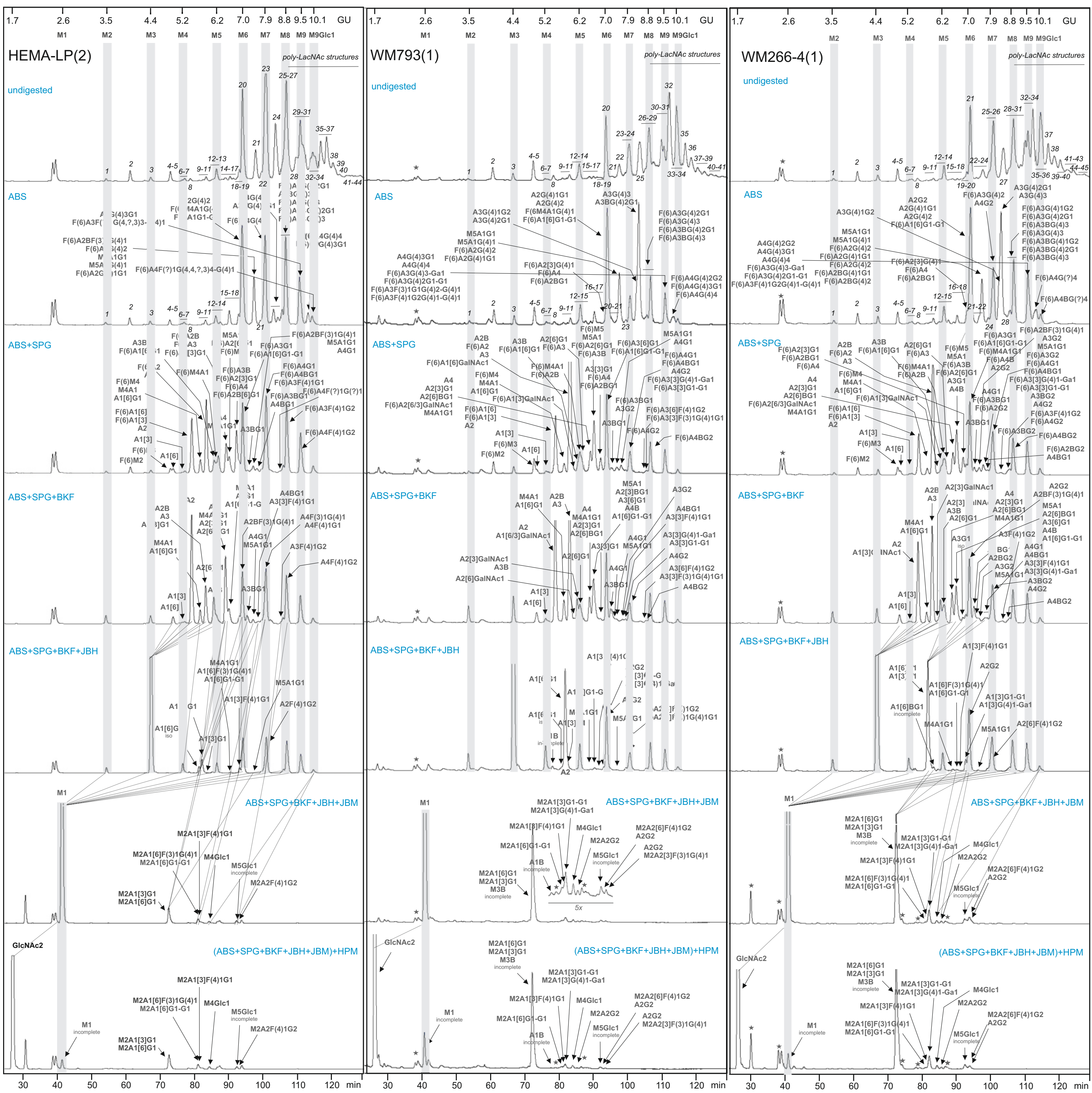

### Supplementary Figures S2-S4.pdf

#### Supplementary Figure S2

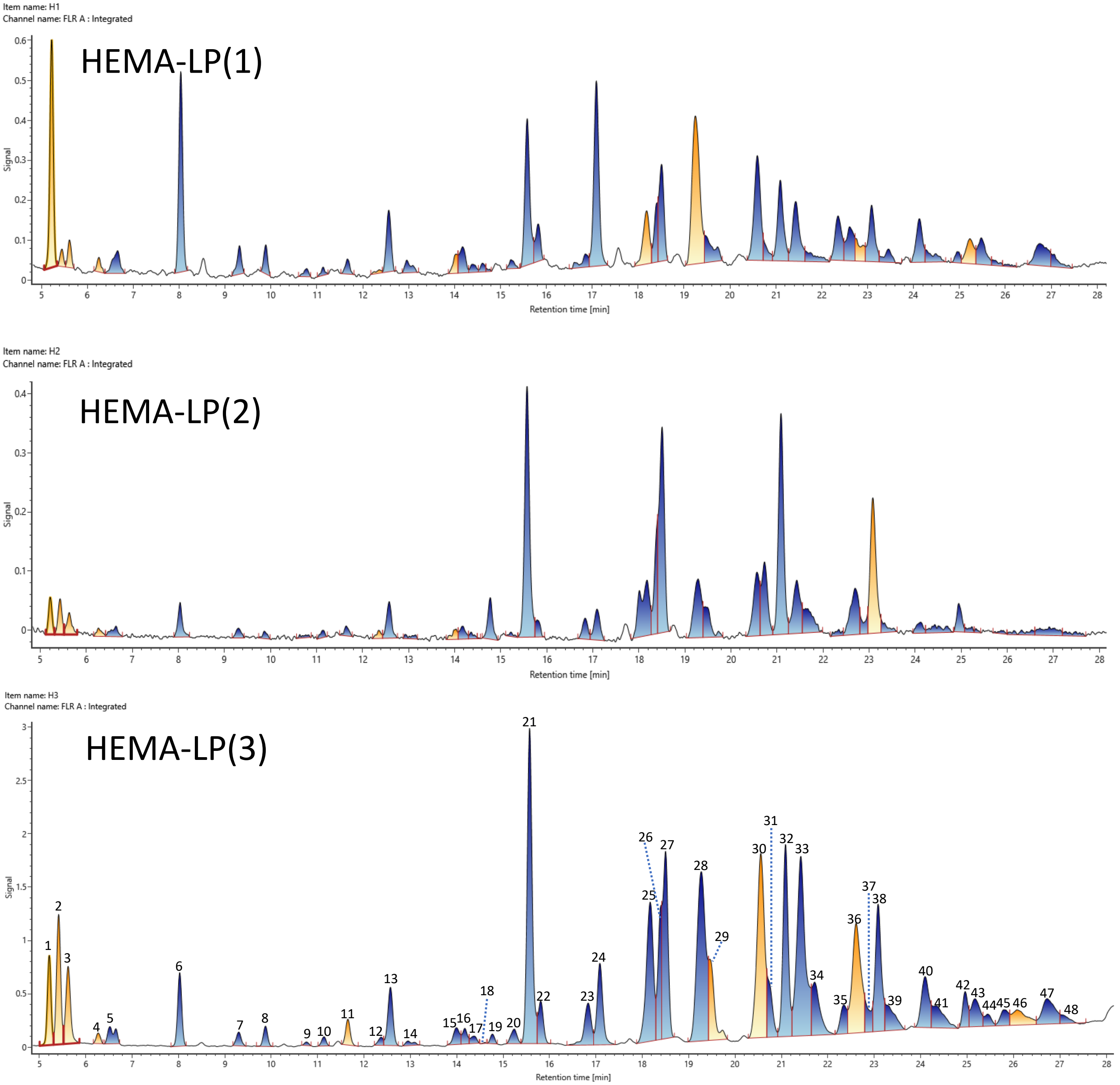

#### Supplementary Figure S3

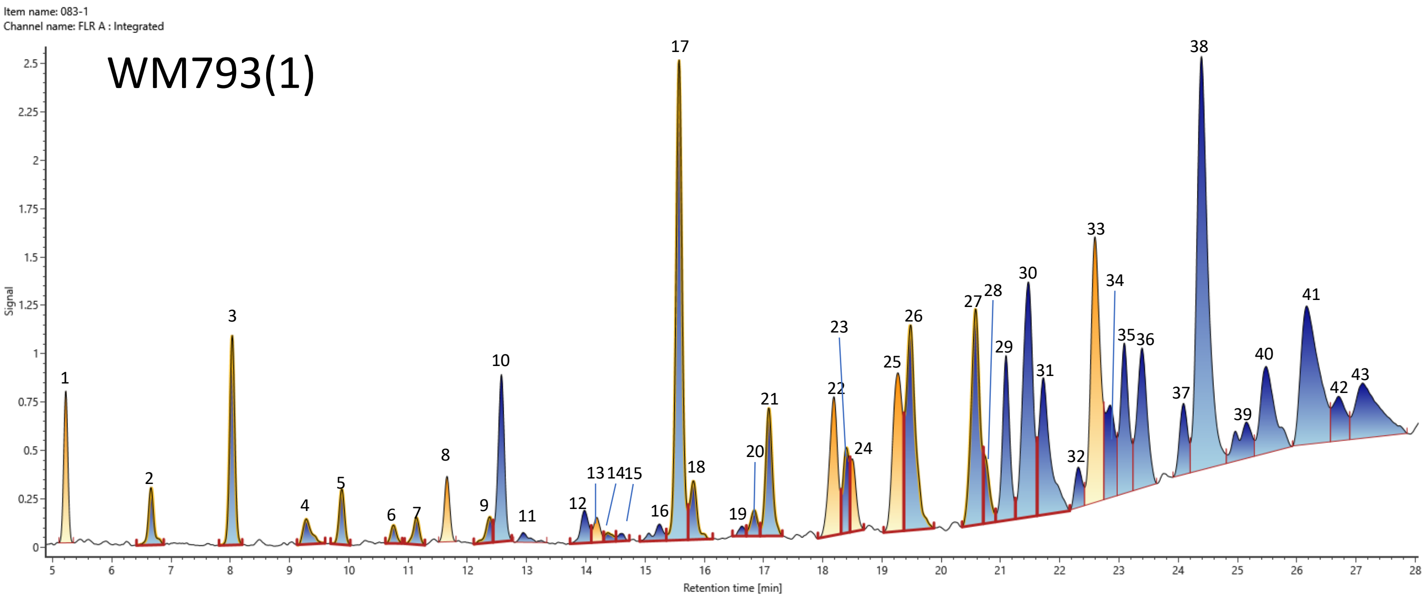


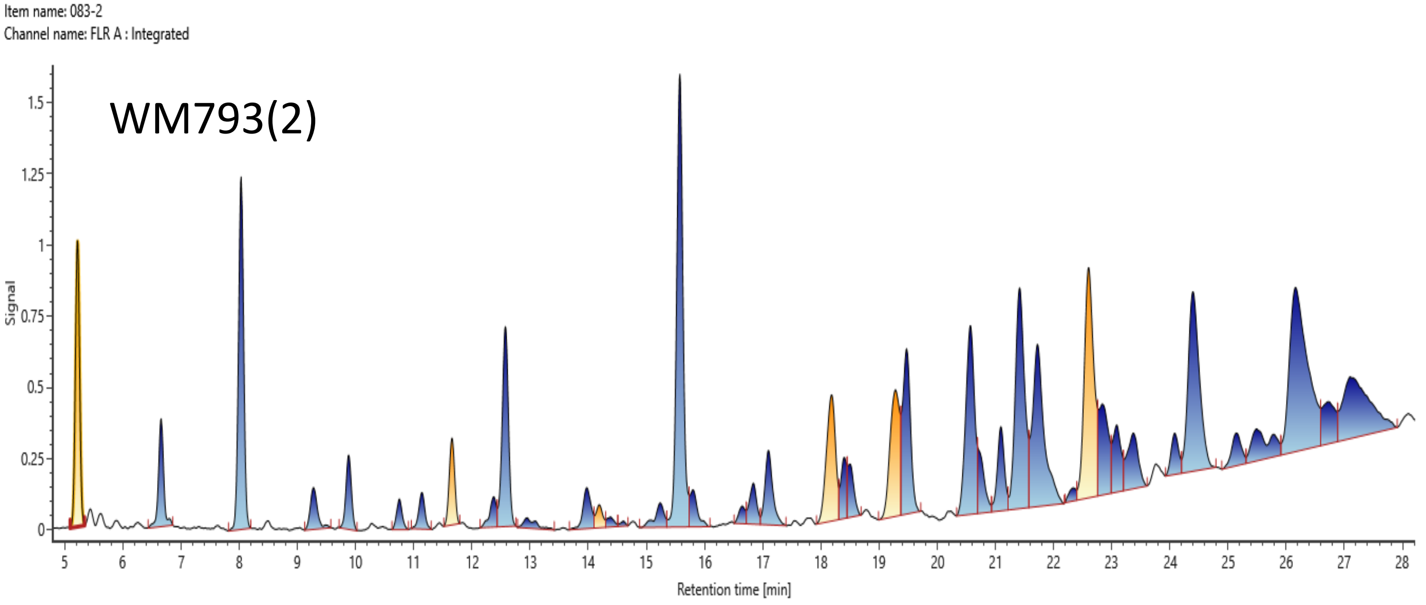


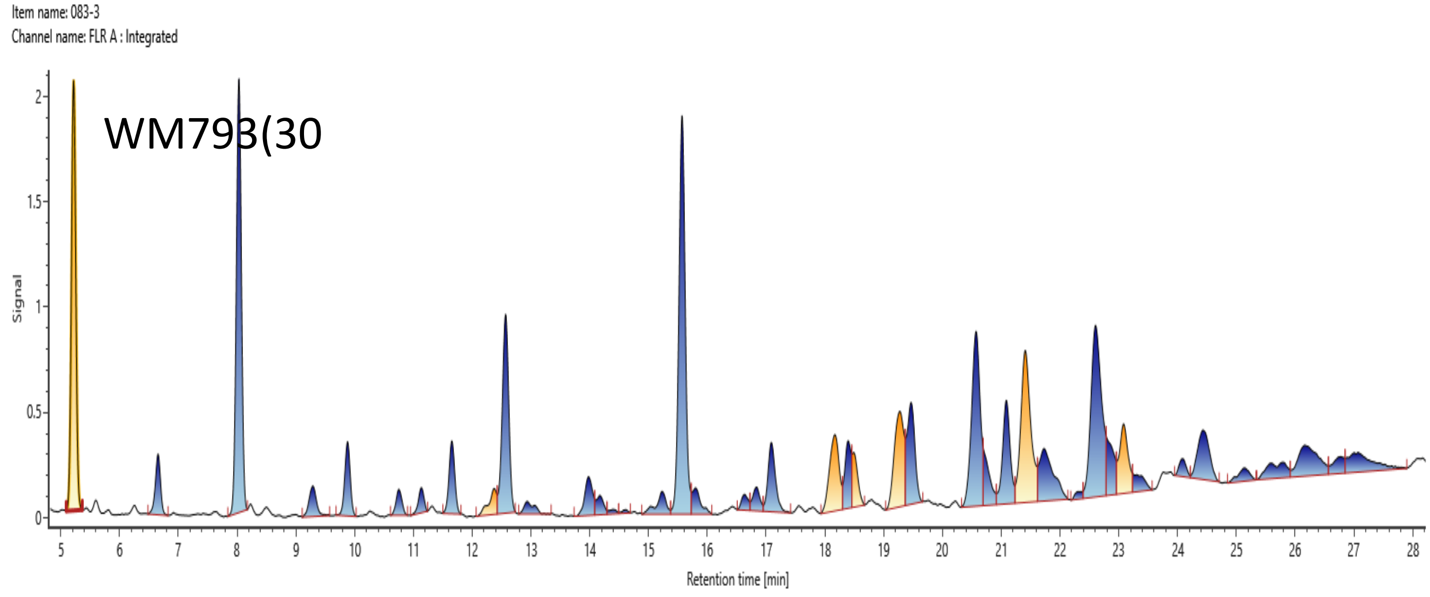

#### Supplementary Figure S3

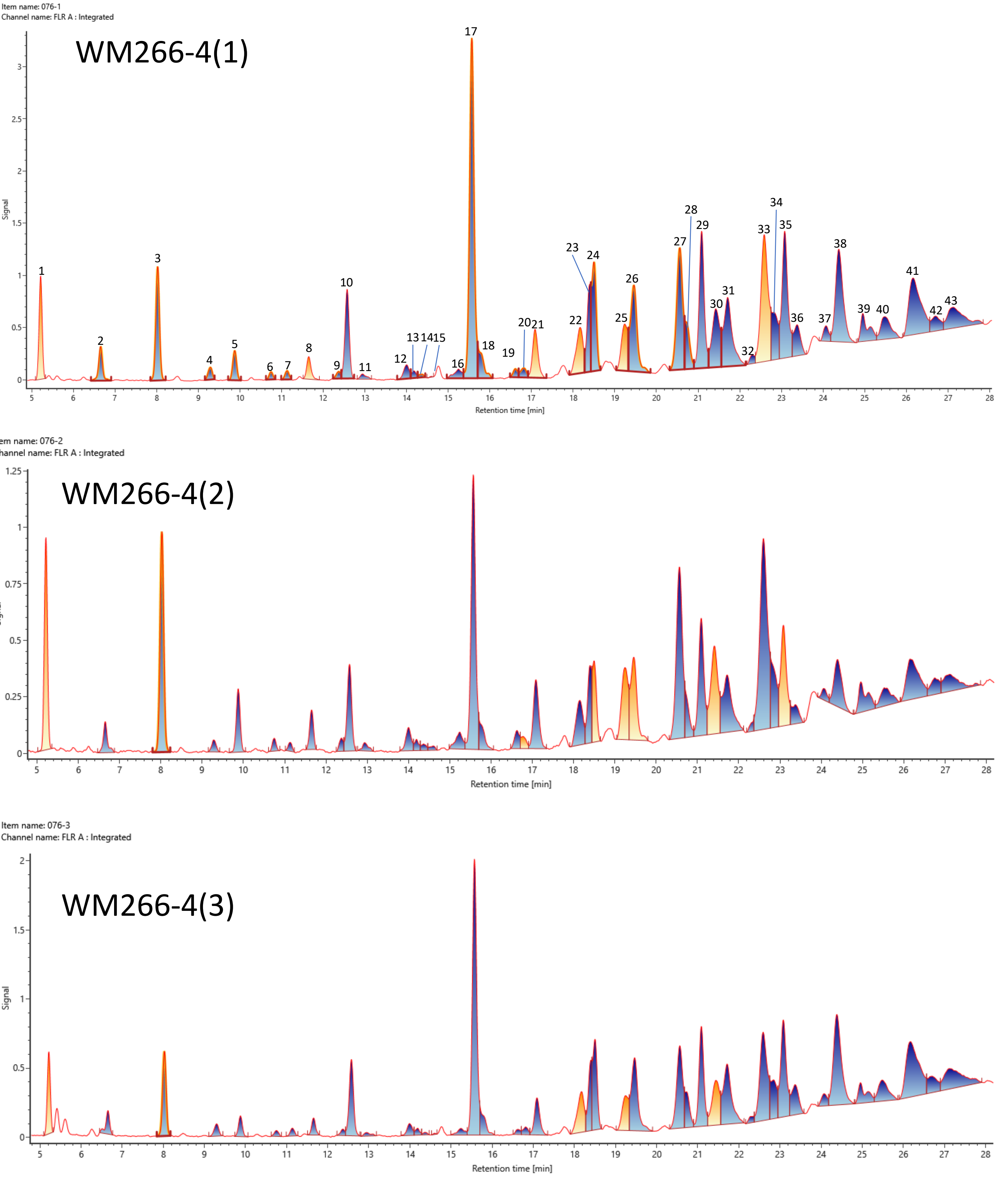

### Supplementary Figures S5.pdf

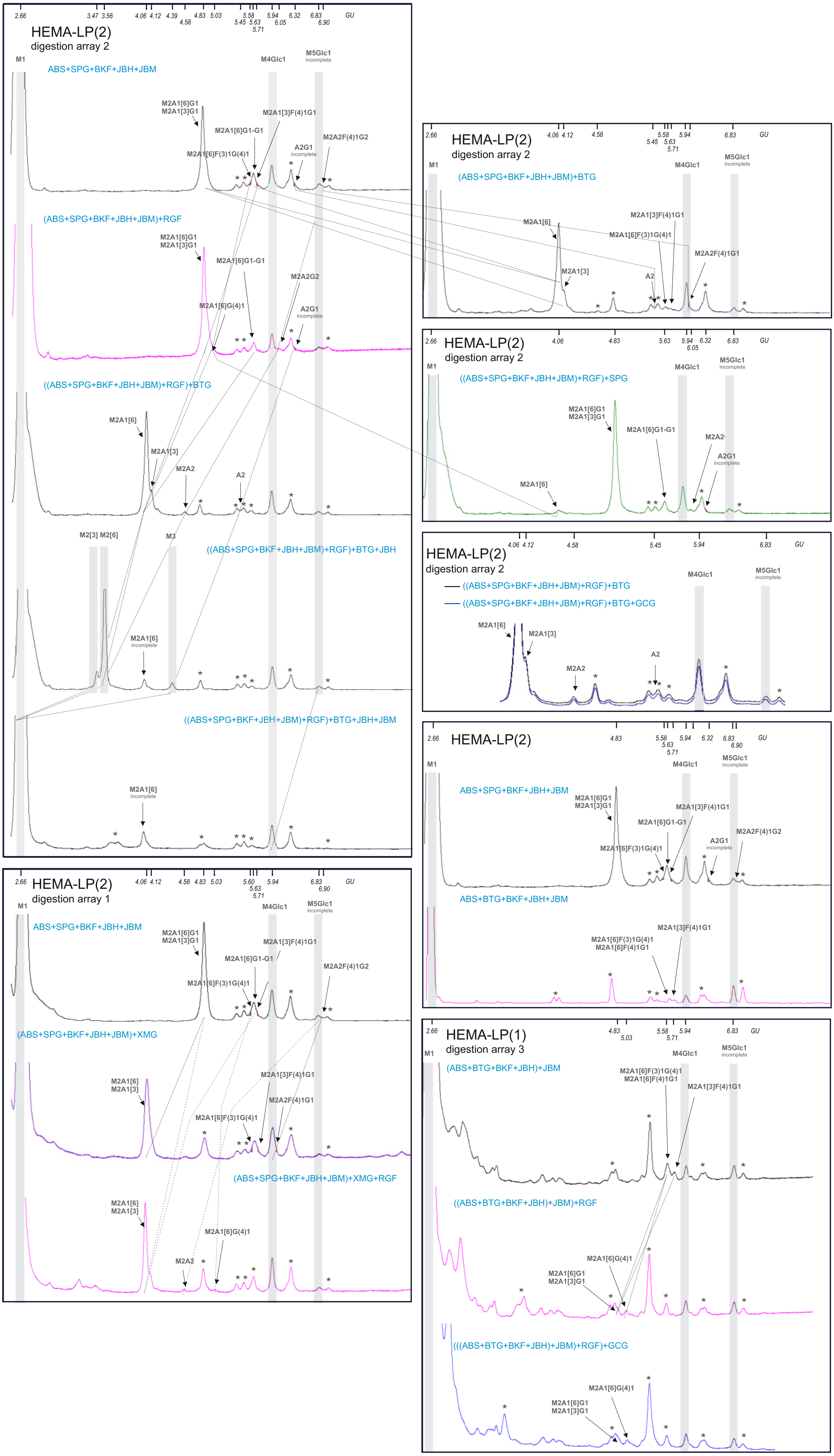

### Supplementary Figures S6.pdf

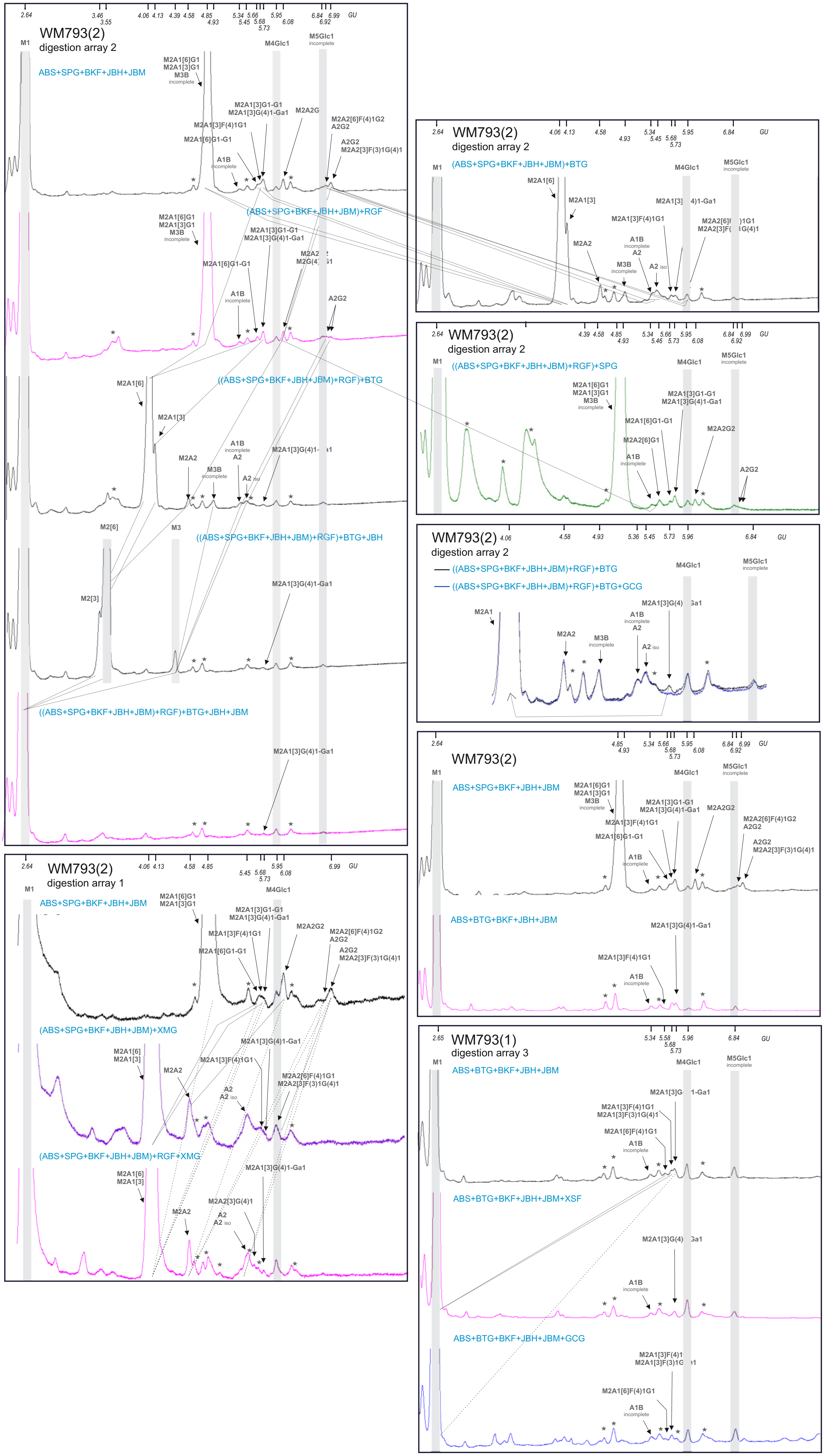

### Supplementary Figures S7.pdf

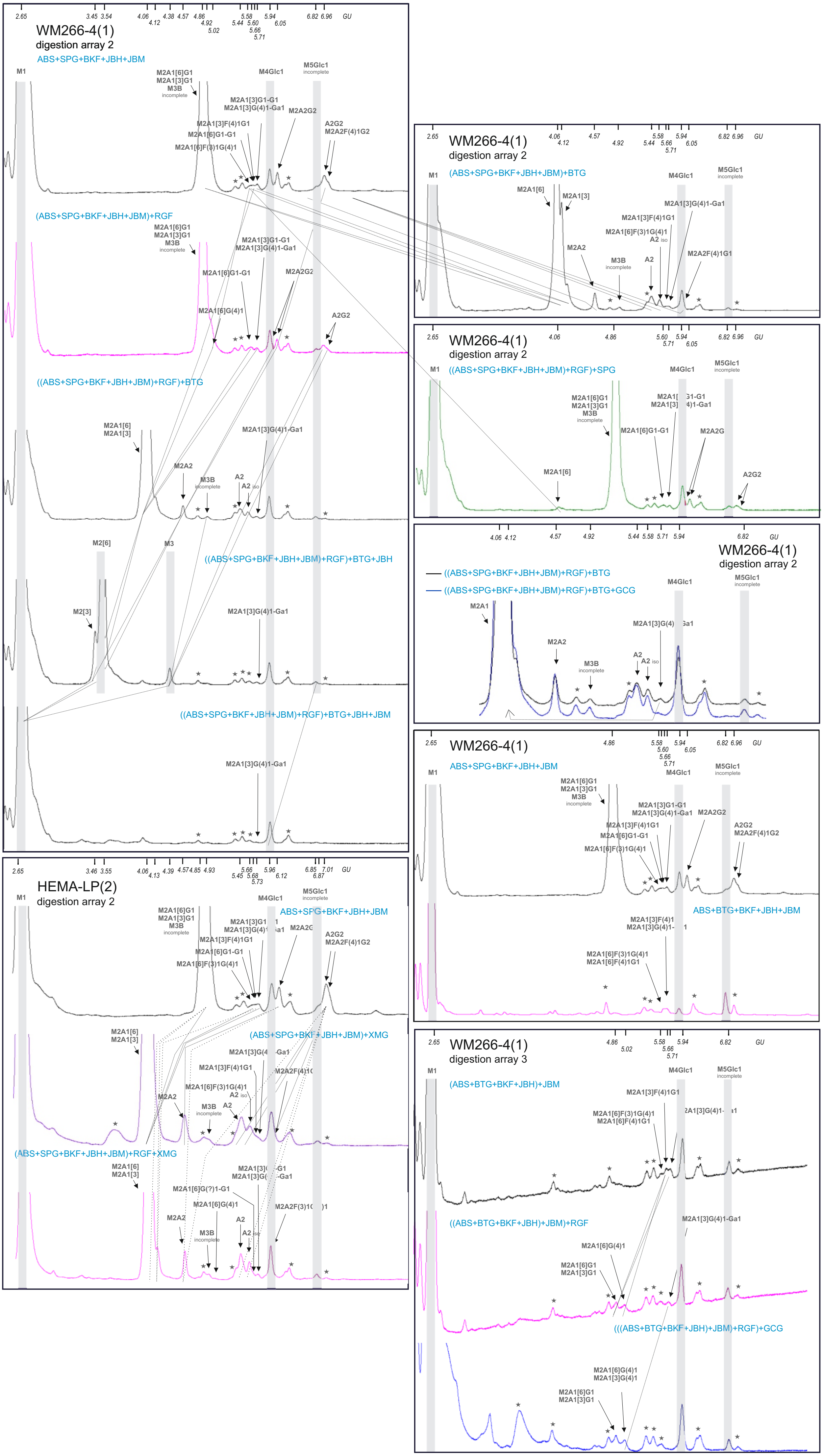

### Supplementary Figures S8.pdf

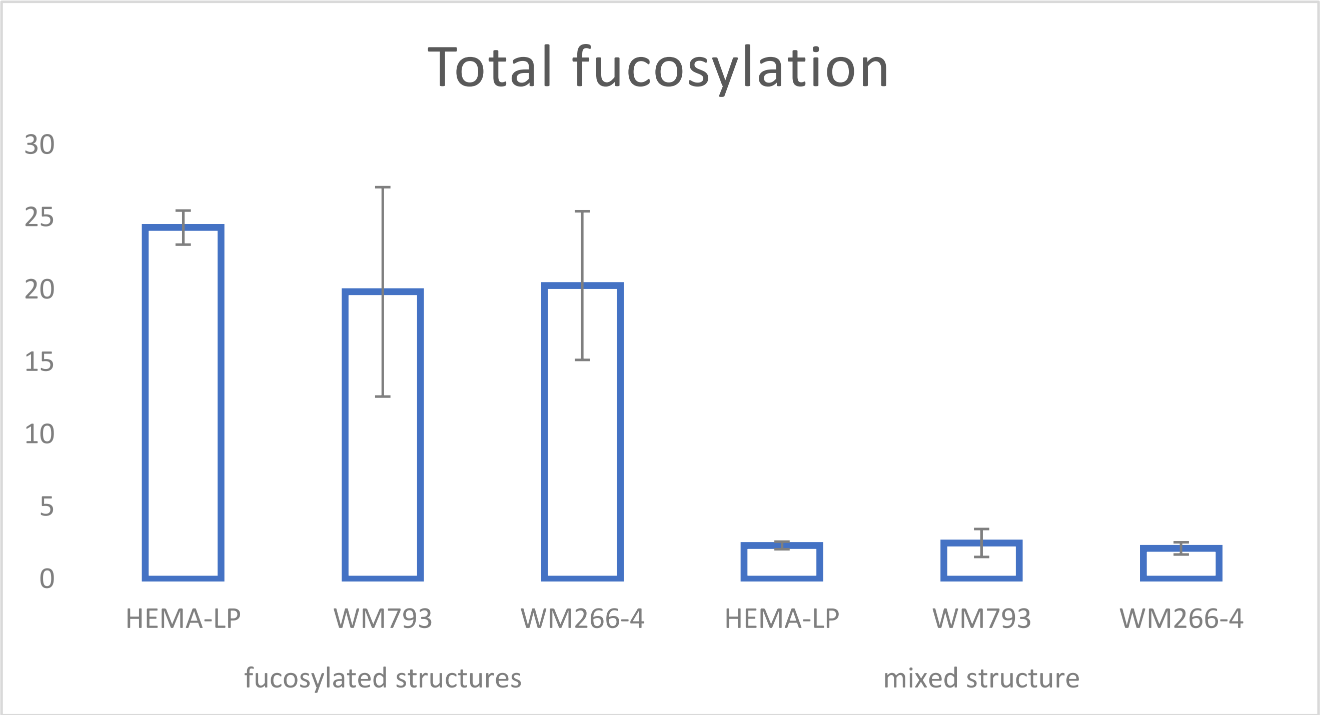
